# Cell Type-specific Hypothalamic Pathways to Brainstem Drive Context-dependent Strategies in Response to Stressors

**DOI:** 10.1101/2023.03.10.532043

**Authors:** Mehran Ahmadlou, Maria Giannouli, Jacqueline F. M. van Vierbergen, Tom van Leeuwen, Wouter Bloem, Janou H. W. Houba, Maryam Yasamin Shirazi, J. Leonie Cazemier, Robin Haak, Mohit Dubey, Fred de Winter, J. Alexander Heimel

**Affiliations:** Circuits, Structure and Function Group, Netherlands Institute for Neuroscience, Meibergdreef 47, 1105 BA Amsterdam, Netherlands; Sainsbury Wellcome Centre for Neural Circuits and Behaviour, University College London, W1T4AJ London, UK; Department of Axonal Signaling, Netherlands Institute for Neuroscience, Meibergdreef 47, 1105 BA Amsterdam, Netherlands; Laboratory for Neuroregeneration, Netherlands Institute for Neuroscience, Meibergdreef 47, 1105 BA Amsterdam, Netherlands

## Abstract

Adaptive behavioral responses to stressors are critical for survival. However, which brain areas orchestrate switching the appropriate stress responses to distinct contexts is an open question. This study aimed to identify the cell type-specific brain circuitry governing the selection of distinct behavioral strategies in response to stressors. Through novel mouse behavior paradigms, we observed distinct stressor-evoked behaviors in two psycho-spatially distinct contexts, characterized by stressors inside or outside the safe zone. The identification of brain regions activated in both conditions revealed the involvement of the dorsomedial hypothalamus (DMH). Further investigation using optogenetics, chemogenetics, and photometry uncovered that glutamatergic projections from the DMH to periaqueductal gray (PAG) mediated responses to inside-stressors, while GABAergic projections, particularly from tachykinin1-expressing neurons, played a crucial role in coping with outside-stressors. These findings elucidate the role of cell type-specific circuitry from the DMH to the PAG in shaping behavioral strategies in response to stressors. These findings have the potential to advance our understanding of fundamental neurobiological processes and inform the development of novel approaches for managing context-dependent and anxiety-associated pathological conditions such as agoraphobia and claustrophobia.

## INTRODUCTION

Adaptive behavioral responses to stressors are critical for survival. Stressors evoke different responses depending on the context in which they are experienced. One way to differentiate these contexts is through a psycho-spatial perspective. This distinguishes stressors that occur outside our perceived safe zone from those that arise within it. Outside stressors, encountered beyond the boundaries of our control, can lead to emotion-focused or passive coping behaviors^1^. In contrast, inside stressors, which are imminent and occur within our safe zone, prompt an active escape-seeking response^2^. Excessive responses to stressors in these two contexts occur in two closely related and relatively common space-associated stress disorders, i.e. agoraphobia^3,4^ and claustrophobia^5,6^. These anxiety disorders reduce the quality of life, impair work performance and carry economic costs^7^. Understanding the neural mechanism responsible for converting threats and their context into behavior is necessary for developing targeted therapies. The concept of psycho-spatial distance of a stressor is closely related to the concept of imminence or proximity of a threat. It is well established that distinct parts of the periaqueductal gray (PAG) in the midbrain are involved in different responses depending on the threat imminence^8^. However, it is an outstanding question which regions of the brain orchestrate the switch between defensive responses depending on the threat imminence^8,9^. One candidate region is the hypothalamus. As the brain controller of the hypothalamic-pituitary-adrenal (HPA) axis it plays a key role in stress and fear responses^7,10,11^. It sends output to the PAG and is hypothesized to contain distinct, but intertwined, defensive systems related to escapable and inescapable threats^12^. The hypothalamus is composed of distinct longitudinal zones^13^. The medial zones contain a number of areas implicated in responses to environmental threats and defensive behavior and are collectively named the medial hypothalamic defensive system^13,14^. Included areas are the anterior hypothalamic nucleus, the dorsomedial portion of the ventromedial hypothalamus and the dorsal premammillary nucleus. The dorsal medial hypothalamus (DMH) lying between these, however, is also activated by stressors (e.g. footshock, thermal stress and exposure to a predator)^13,15,16^ and blockade of GABAA receptors in this area causes well-coordinated escapes^17^. Previously, it was shown that different areas within the medial hypothalamic defensive system can process different threats^11,18^. It has remained unclear whether cell classes within regions of the hypothalamus are differentially involved in expressing stress responses depending on the psycho-spatial context.

Here, we first developed behavioral tests to investigate behavioral correlates of mice exposed to the two conditions, stressors inside and outside the safe zone. Using optogenetics and calcium fiber photometry, we showed that two separate populations of neurons in the DMH are essential for these two contextually different conditions which consequently lead to two different behavioral responses. Finally, we showed that for both behavioral responses the projection of these two neural populations to the key region in the midbrain fear circuit, i.e. the periaqueductal gray^19^, is required.

## RESULTS

### Behavioral responses to inside- and outside-stressors

First, we designed a paradigm where we provide stressors in close proximity to the mouse and allow the mouse a potential escape-route. In this paradigm, the mouse is in a plexiglass box (with an open top) for ten minutes, where all safe places (i.e. the four corners) have cotton balls with an innately threatful odor of a predator, 2,5-dihydro-2,4,5-trimethylthiazoline (TMT)^20^ (**Figure 1A, Video S1**). The results showed a dramatically larger number of jumps compared to the control (without TMT) (**Figure 1B; Figures S1A-F**). The number of jumps when all corners contained TMT cotton balls was also significantly higher than when mice had safe corner opportunities (**Figure 1C, Video S1**). Therefore, this paradigm provides a situation where the stressors are placed inside the safe zone and provoke robust escape-seeking behavior.

**Figure 1.**
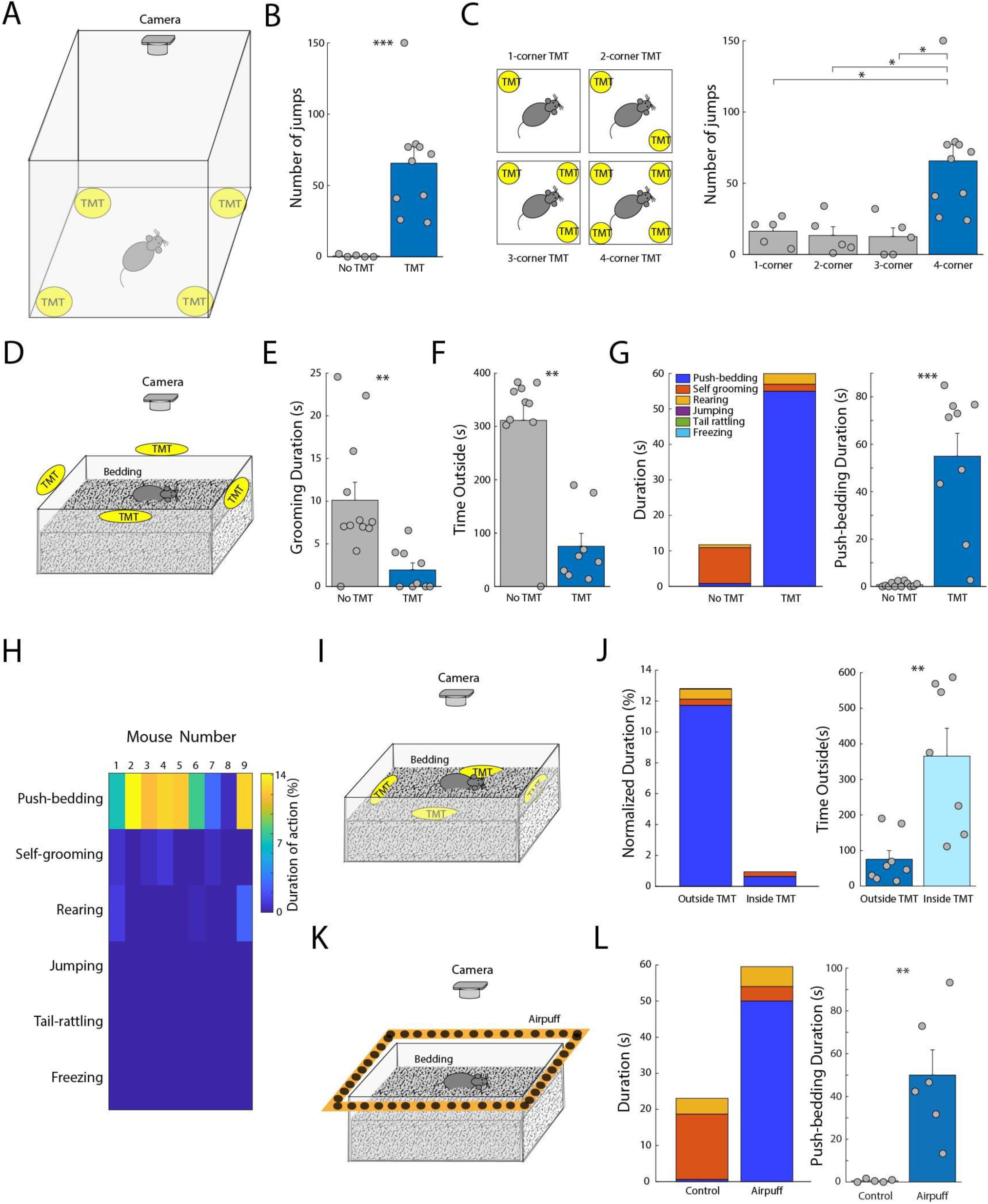
Behavioral fingerprints for “being in an inside-stressor situation” and “being trapped in an outside-stressor situation”. (**A**) Schematics of an inside-stressor test with TMT cottons located at all corners. (**B**) Bar graph shows the number of jumps taken by C57BL/6 mice in the inside-stressor test without (N = 5 mice) and with TMT (N = 10 mice). (**C**) Schematics of the inside-stressor test with TMT cottons located at 1, 2, 3 and 4 corners. Bar graph shows the number of jumps taken by C57BL/6 mice in the inside-stressor test with 1, 2, 3 and 4 corners filled with TMT cotton (N = 5, 5, 5 and 10 mice, respectively). (**D**) Schematics of an outside-stressor test with TMT cottons sticking at all edges outside the home-like cage with a thick layer of beddings. (**E**) Bar graph shows duration of self-grooming without (N = 12 mice) and with (N = 9 mice) TMT. (**F**) Bar graph of duration of time spent outside the cage (defined by all paws and body lifted up from the bedding surface) without and with TMT in C57BL/6 mice (N = 10 and 8 mice, respectively). (**G**) Stacked bar graph shows duration for each action (bedding-pushing, self-grooming, rearing, jumping, tail rattling and freezing) in the outside-stressor test (without and with TMT) taken by the mice in (E). The bar graph shows duration of bedding-pushing in the outside-stressor test without and with TMT taken by the mice in (E). (**H**) Heatmap shows the fraction of time that individual mice in (G) spent in each behavior in presence of TMT. (**I**) Schematics of an inside-stressor test with TMT cottons located at all corners inside the home-like cage with a thick layer of bedding. (**J**) Stacked bar graph shows duration of the actions (bedding-pushing, self-grooming, rearing, jumping, tail rattling and freezing) when the TMT cottons are outside or inside the cage, normalized by the duration of time spent inside the cage. The bar graph shows the duration of time spent outside the cage when the TMT cottons are outside or inside the cage (N = 8 and 7 mice, respectively). (**K**) Schematics of an outside-stressor test with air puff holes located at all edges outside the home-like cage with a thick layer of bedding. In the 10-min period prior to the test, mice receive air puffs upon sniffing the air puff holes. The behavioral response is measured in the next 10 minutes without the exposure to the air puff. (**L**) Stacked bar graph shows duration of the actions (bedding-pushing, self-grooming, rearing, jumping, tail rattling and freezing) without (control) and with air puff prior to the test. The bar graph shows the duration of bedding-pushing in both conditions (N = 5 and 6 mice, respectively). *: p < 0.05, **: p < 0.01, ***: p < 0.001. See also Figure S1, Video S1 and Table S1.

The behavior of mice being trapped in a situation where the threat comes from the outside (psycho-spatially distal threat) is yet unclear. In order to find out how mice behave when they are in such a situation, we made a paradigm in which the mouse is in an open cage with a thick layer of bedding (to provide an environment similar to its home cage) and we placed cotton balls with TMT directly surrounding the cage (**Figure 1D**). The mouse could easily leave the cage. We measured well-studied defensive behaviors (jumping, freezing, tail-rattling), self-grooming, rearing and bedding-pushing. The latter is the displacement of the bedding by, mostly, the snout and partially by forepaws (**Video S1**). The movements are similar to defensive burying^21^, but without an object to bury. After a short period of exploration (rearing, sniffing the cotton balls), mice sensed the outside-stressor. They showed less grooming and spent much more time inside the cage, compared to the control group where no TMT was placed outside the cage (**Figures 1E and 1F**). Interestingly, mice showed significantly higher duration of bedding-pushing compared to the control group and no significant change in other defensive behaviors (**Figures 1G and 1H**). The more intense the stressor was, the higher was the duration of the bedding-pushing behavior (**Figures S1G-I**). Moreover, to see whether the bedding-pushing is a unique coping behavior specific to being trapped by an outside-stressor or whether it is also evoked in an inside-stressor situation, we placed the TMT cottons inside the cage with bedding (**Figure 1I**). In this situation, mice did not show bedding-pushing and they preferred to escape the box (**Figure 1J**). To test whether the bedding-pushing behavior occurs only in response to TMT or olfactory stressors and not to other stressors, we trapped the mouse by installing air puffers around the box. Whenever the mouse investigated the possibility to venture out of the cage, it received an air puff (**Figure 1K, Video S1**). After ten minutes in which the mouse learned of the existence of this outside-stressor, we scored its behavior for ten minutes. Interestingly, again among all fear responses, mice predominantly showed bedding-pushing behavior and did so for much longer than control animals surrounded by inactive puffers (**Figure 1L**). This suggests that the bedding-pushing is not a response that is specific to predator smell but to an outside-stressor in general. In addition, we aimed to differentiate the possible confounding effect of the varied environments utilized in the inside-stressor test (without beddings in the small plexiglass box) from the outside-stressor test (specifically, the presence of beddings), in relation to the stressor location and its impact on the observed behavioral responses. To this end, we video recorded the behaviors when mice were in the inside-stressor test (with TMT), but with bedding inside the box (**Figure S1J**). Consistent with our results (**Figure 1**), mice showed a significant increase in the number of jumps, but no significant increase in the push-bedding behavior (**Figures S1K and S1L**).

### Excitatory neurons in DMH are required for inside-stressor fear response

To determine where the distinction is made in the two responses to the stressors based on their spatial location, we investigated which regions within the hypothalamus are activated in both conditions. The activity marker cFos appeared shortly after both stress conditions in a few subregions of the hypothalamus (**Figures 2A-2C and S2**). Interestingly, one of these was the dorsomedial hypothalamic region (DMH), a region previously implicated in anxiety, stress and fear^13,15–17,22–24^. Recent studies have shown that pharmacologically activating DMH induces defensive alertness behavior and vertical oriented jumping^25^, which is distinct from the well-studied startle and “explosive” defensive behavior and jumping induced by activating the midbrain defensive circuit (e.g. PAG)^19,25,26^. However, DMH was not previously implicated in the two psycho-spatially distinct fear responses, i.e. escape jumping instigated by the inside-stressor situation and the bedding-pushing coping behavior in response to the outside-stressor situation.

**Figure 2.**
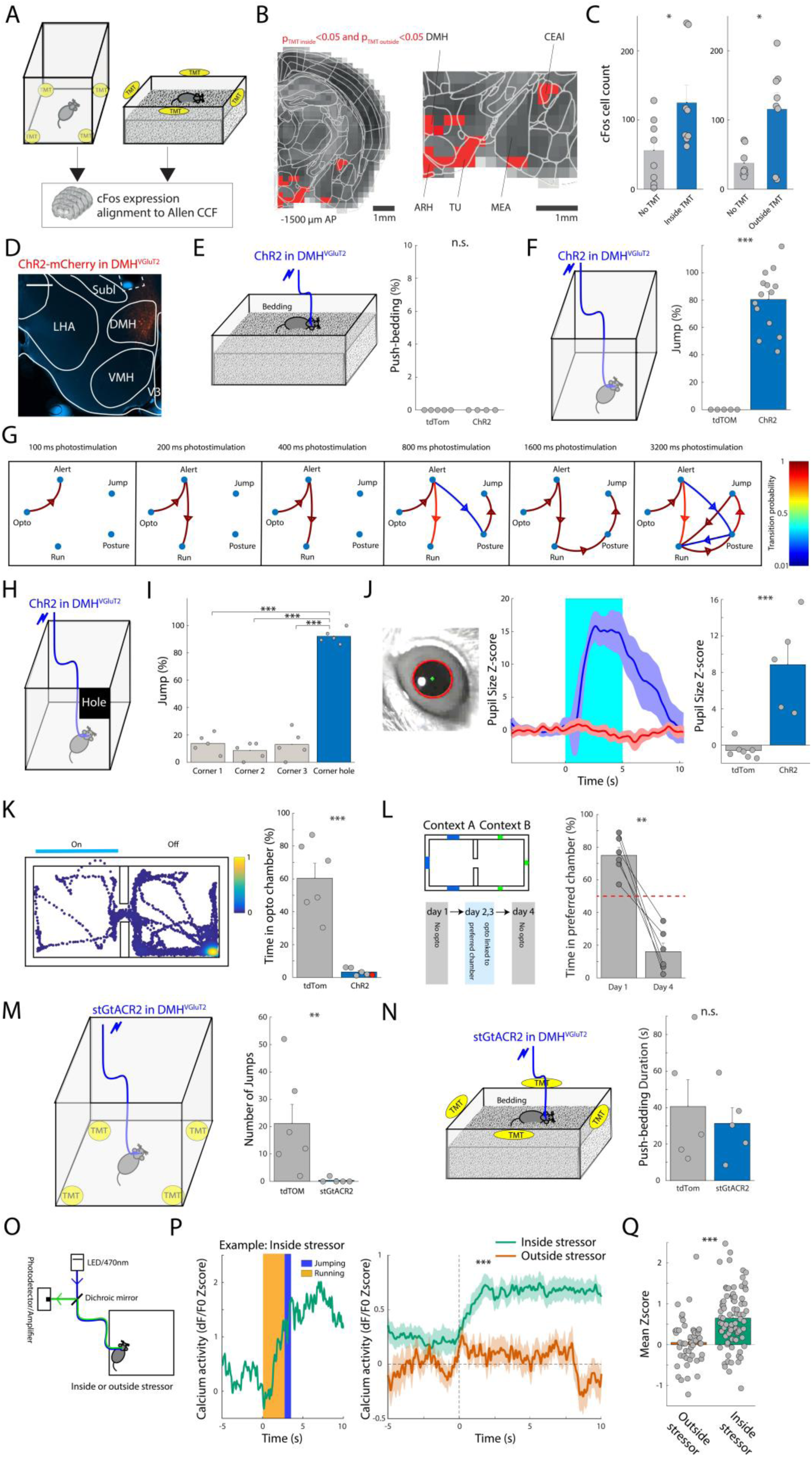
Excitatory neurons in DMH are required for inside-stressor fear response. (**A**) Mice were either exposed to the inside-stressor or outside-stressor before immunohistochemistry for cFos expression in about 30 coronal slices (1.5mm) including the hypothalamus, which were subsequently aligned to the Allen Common Coordinate Framework. (**B**) Red squares indicate the voxels at 1.5 posterior to Bregma that had significantly more (t-test, p<0.05) cFos positive neurons for both the inside-stressors and outside-stressors paradigms. The left and right hemispheres were overlaid. Voxel size is 0.25 mm. Several areas including significant voxels are indicated, DMH: Dorsomedial nucleus of hypothalamus, CEAl: Central amygdala nucleus, lateral part, ARH: Arcuate hypothalamic nucleus, TU: Tuberal nucleus, MEA: Medial amygdala nucleus. (**C**) The counts for cFos positive cells in the red DMH voxel indicated by the blue outline in the right part of (C) are higher for both the inside-stressor and outside-stressor compared to the controls without TMT. (**D**) Expression of AAV-ChR2-mCherry in DMH of a VGluT2-Cre mouse. Scale bar indicates 0.25 mm. Dashed line indicates the fiber position. DMH: Dorsomedial nucleus of hypothalamus, LHA: Lateral hypothalamic area, SubI: subincertus nucleus, VMH: Ventromedial nucleus of hypothalamus, V3: Third ventricle. (**E**) Schematics of photo stimulation of DMH^VGluT2^ neurons in the home-like cage with beddings (without TMT). The bar graph shows the number of bedding-pushing occurrences by each photo stimulation normalized to the number of photo stimulation trials (in percentage) in tdTomato and ChR2 injected mice (N = 5 and 4 mice, respectively). (**F**) Schematics of photo stimulation of DMH^VGluT2^ neurons in the box (without TMT). The bar graph shows the number of jumps by each photo stimulation normalized to the number of photo stimulation trials (in percentage) in tdTomato and ChR2 injected mice (N = 5 and 14 mice, respectively). (**G**) Transition probability between the measured actions (alertness (sudden head/body orienting), run, posture (posture preparation for jumping), jump and the onset of photo stimulation of the DMH^VGluT2^ neurons with different durations of photo stimulation (N = 20 trials per photo stimulation duration, 3 mice). The color bar indicates probabilities between 0.01 and 1. Transition probability less than 0.01 is shown with no edge between the nodes, for better visualization. (**H**) Schematics of photo stimulation of DMH^VGluT2^ neurons in the box (without TMT) with a hole in one top corner (corner hole). (**I**) The bar graph shows the number of jumps to each of the top corners by each photo stimulation normalized to the number of photo stimulation trials (in percentage) in ChR2 injected mice (N = 5 mice). (**J**) Z-score (middle) and mean Z-score (right) of pupil size of tdTom (red; N = 7 mice) and ChR2 (blue; N = 5 mice) mice with photo stimulation from 0 to 5 s. (**K**) An example heatmap of the track of a ChR2 mouse in a real time place preference/aversion (RTPPA) test. The bar graph shows the duration of time that control tdTom and ChR2 (N = 6 and 6 mice, respectively) mice spent in the opto-linked chamber in the RTPPA test. Red data point indicates the example mouse presented in the heatmap track. (**L**) Schematics of the RTPPA conditioning test, where ChR2 mice on day 2 and day 3 are photo stimulated in one chamber (the preferred chamber in the pre-conditioning day). On day 1, pre-conditioning, and day 4, post-conditioning, there is no photo stimulation. The bar graph shows duration of time spent in the preferred chamber before and after conditioning (N = 6 mice). The red dashed line represents the chance level (50%). The duration after conditioning is significantly lower than chance level (T-test; p-value: 0.0003, t[6] = 7.4488). (**M**) Schematics of photo inhibition of DMH^VGluT2^ neurons in the inside-stressor test with the TMT cottons. The bar graph shows the number of jumps (in the 10-min test) in tdTomato and stGtACR2 injected mice (N = 6 and 5 mice, respectively). (**N**) Schematics of photo inhibition of DMH^VGluT2^ neurons in the home-like cage with beddings with outside-stressor (TMT). The bar graph shows the duration of bedding-pushing (in the 10-min test) in tdTomato and stGtACR2 injected mice (N = 5 and 5 mice, respectively). (**O**) Schematic of calcium photometry of DMH^VGluT2^ neurons in inside- and outside-stressor tests. (**P**) Left: Example Calcium activity of DMH^VGluT2^ neurons (dF/F0 Z-score) in an inside-stressor test. Yellow and blue bars indicate duration of running and jumping. Right: Calcium activity of DMH^VGluT2^ neurons (dF/F0 Z-score) in inside-(green) and outside-stressor (orange) tests, averaged over trials. The light color around the average indicates the standard error. Time 0 s in inside-stressor test is the onset of running away (or running + jumping) and in outside-stressor test is the onset of bedding-pushing. (**Q**) Mean Z-score of calcium activity in response to stressors in outside- and inside-stressor tests (N = 47 and 82 events, 4 mice, respectively). *: p < 0.05, **: p < 0.01, ***: p < 0.001. See also Figures S2 and S3, Video S2 and Table S1.

Next, we questioned whether the DMH is involved differently in these two inside- and outside-stressor situations. One possibility is that activity signaling of fear or stress in general, regardless of the situation, is present in the DMH, and that the differences in behavioral outcome depend on different inputs or processing in downstream areas. Another possibility is that the neural activity is already situation-specific in the DMH. The hypothalamus harbors many cell-types, but the major division is into excitatory glutamatergic neurons and GABAergic inhibitory neurons^27^. As the downstream effect of these neurons is likely to be very different, we first manipulated the excitatory neurons in the DMH.

In order to activate the excitatory neurons in the DMH, we expressed ChR2 bilaterally in DMH of VGluT2-Cre mice through injection of an AAV virus (**Figure 2D**). Upon activation, mice did not show any bedding-pushing when they were in the cage with bedding (**Figure 2E**), but they showed a dramatic increase in the number of jumping in the open field (**Figure 2F**), and in all conditions: in the absence of bedding in the open field (**Video S2**), in the presence of bedding in the open field (**Video S2**), or in the presence of bedding in the cage (**Video S2**). They showed no other types of defensive behavior (no freezing, no tail rattling). Next, we did photostimulation of different durations. This showed that the induced jumping was not a startle response, but it was a coordinated jump evidenced by a progressive behavior over time (initiated by orienting the head/body, followed by running, making a jumping posture and ending by jumping) (**Figure 2G, Video S2**). We examined whether the induced jumping is goal-directed, by activating the DMH^VGluT2^ neurons in an open field box with an opening at one of the top corners (**Figure 2H**). Interestingly, the jumps induced by the photostimulation were tremendously goal-directed towards the top open corner (**Figure 2I**), and the preference for the corner with the top open was not merely because the mouse was close to that corner when the light was not yet activated (**Figure S3A**). These results imply that the photostimulated jumping is a top-down goal-directed behavior which needs integrative processing, rather than a bottom-up startle response.

Furthermore, to test whether the observed jumping was an escape behavior or purely a locomotive behavior, we examined whether the photostimulation induces aversion and negative arousal. Therefore, we optogenetically activated DMH^VGluT2^ neurons in head-fixed mice and video recorded the pupil and whiskers. The pupil size and whisker activity measures showed an increased arousal level by activation of the DMH^VGluT2^ neurons (**Figures 2J and S3B**). The valence was examined using a real-time place preference/avoidance test in a double-chamber, where one of the chambers is linked to the optogenetic light (light-chamber). Compared to control mice which were injected with a tdTomato-vector, activation of the DMH^VGluT2^ neurons significantly decreased the time spent in the light-chamber (**Figure 2K**). Moreover, one day after two sessions of conditional activation of the DMH^VGluT2^ neurons in the real-time place preference/avoidance test, in the absence of photostimulation, mice preferred the chamber in which no activation took place, indicating the formation of an associative fear memory (**Figure 2L**). These results together showed that activation of the DMH^VGluT2^ neurons induces a negative arousal commonly associated with fear and stress. Moreover, photostimulation of the DMH^VGluT2^ neurons in presence of their own nesting material (decreasing stress) or cotton balls with TMT (increasing stress), revealed that the number of the induced jumps was modulated by the stressor levels (**Figures S3C-S3F**).

Next, we questioned whether the DMH^VGluT2^ neurons are necessary for expressing fear of the inside-stressor situation. To answer this, we expressed stGtACR2 in the DMH^VGluT2^ neurons and photo-inhibited them in the inside-TMT paradigm. Interestingly, compared to the tdTomato control mice that also had an optic fiber connected, the jumping behavior was dramatically suppressed (**Figure 2M**) and there were no other significant stress responses (freezing and tail rattling). Furthermore, photo-inhibition of the DMH^VGluT2^ neurons did not significantly suppress the bedding-pushing behavior in the outside-stressor paradigm (**Figure 2N**). Moreover, calcium photometry of GCaMP6s expressed in DMH^VGluT2^ neurons (**Figure 2O**) showed that these neurons were activated during running/jumping induced by stressors in the inside-stressor situation, but not by bedding-pushing behavior in the outside-stressor situation (**Figures 2P and 2Q**). Notably, the increase of activity in the inside-stressor situation started from onset of running and before jumping (**Figure S3G**). Therefore, activity of DMH^VGluT2^ neurons is specific for the fear of an inside-stressor and the ensuing response and not for the fear of an outside-stressor and the corresponding response.

### Inhibitory neurons in DMH are required for outside-stressor fear response

It has been shown that DMH is important in the regulation of acute hypertensive and tachycardic responses to mental stress in an air-jet stress paradigm^28,29^. As DMH^VGluT2^ neurons did not influence the fear response to outside-stressor, we asked whether another population of neurons, rather than the excitatory neurons in the DMH plays a role in the outside-stressor fear response. To this end, we targeted the inhibitory neurons in the DMH. In order to activate the inhibitory neurons in the DMH, we expressed ChR2 bilaterally in DMH of GAD2-Cre mice through injection of an AAV virus (**Figure 3A**). Activation of DMH^GAD2^ neurons did not cause any jumping in the open field (**Figure 3B**), but dramatically increased bedding-pushing when the mice were in the cage with bedding (**Figure 3C**), while there was no significant change in other defensive behaviors (freezing, running, jumping and tail rattling). Presence of bedding was essential for the expression of the behavior. More than VGluT2, GAD2 is expressed in many neurons in nearby brain regions, and there is a risk of the virus spreading into these regions. Knowing that Tachykinin 1 (TAC1), which encodes an important neuropeptide for stress regulation and anxiety-related behavior^30,31^, substance P, is expressed in DMH (https://portal.brain-map.org/,^32^) and that substance P in DMH plays a key role in controlling physiological responses to stress, through the HPA axis^33^, together with the evidence that in some thalamic and subthalamic regions (e.g. thalamic reticular nucleus, lateral geniculate nucleus and zona incerta) TAC1 is exclusively expressed in inhibitory neurons^34,35^ (https://portal.brain-map.org/), we examined whether TAC1 is also expressed in the DMH inhibitory neurons. Our double-color in-situ hybridization and cre-line characterizations^36^ showed that DMH^TAC1^ neurons are inhibitory (**Figures 3D and S4A**). Activation of the DMH^TAC1^ neurons (**Figure 3E**) showed the same results as activation of the encompassing group of inhibitory neurons, i.e. exclusively inducing bedding-pushing behavior (**Figure 3F, Video S3**) and absolutely no jumping behavior. The same results were observed in the plexiglass box.

**Figure 3.**
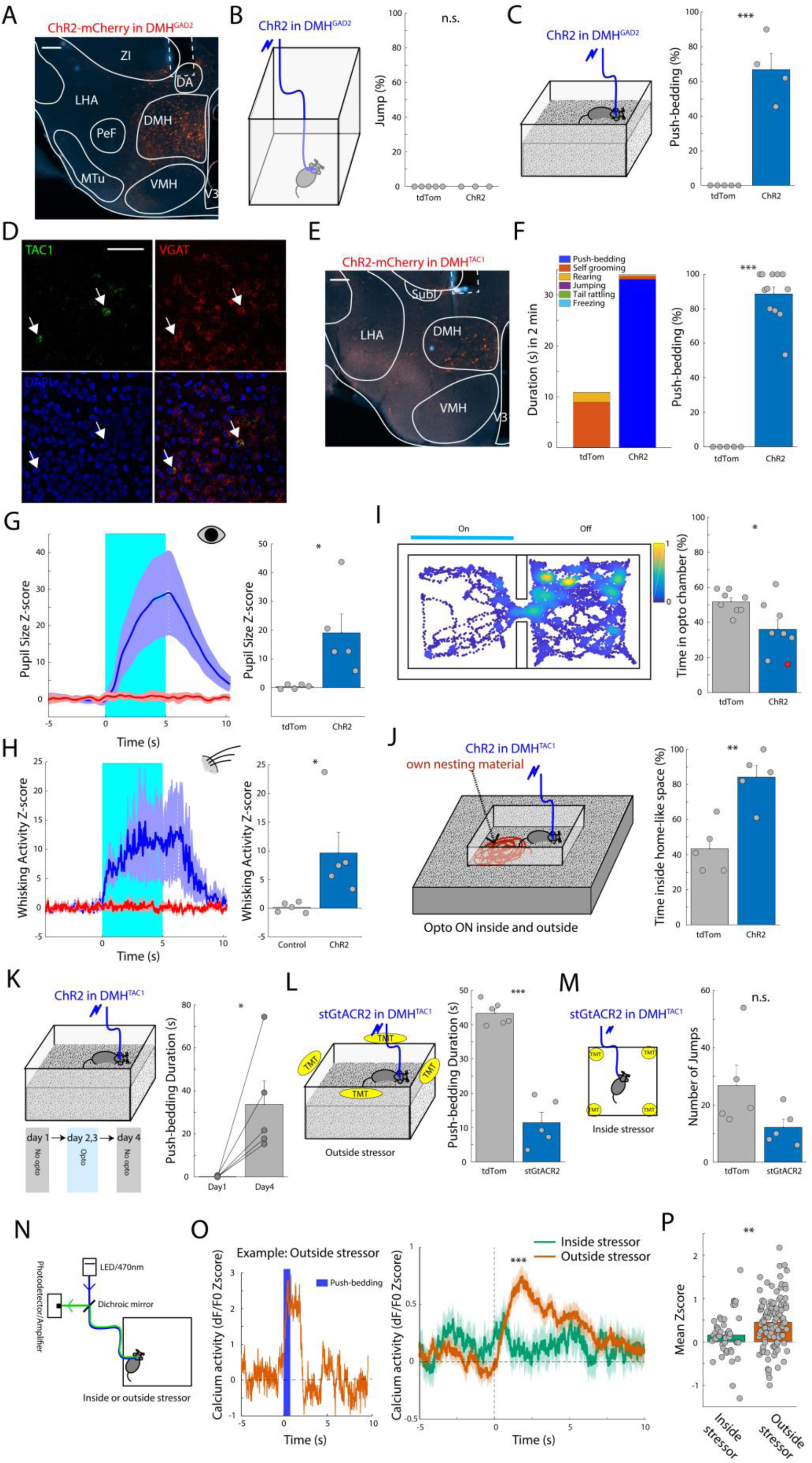
Inhibitory neurons in DMH are required for outside-stressor fear response. (**A**) Expression of AAV-ChR2-mCherry in DMH of a GAD2-Cre mouse. Scale bar indicates 0.2 mm. Dashed line indicates the fiber position. DA: Dorsal hypothalamic area, DMH: Dorsomedial nucleus of hypothalamus, LHA: Lateral hypothalamic area, MTu: Medial tuberal nucleus, PeF: Perifornical nucleus, VMH: Ventromedial nucleus of hypothalamus, V3: Third ventricle, ZI: Zona Incerta. (**B**) Schematics of photo stimulation of DMH^GAD2^ neurons in the box (without TMT). The bar graph shows the number of jumps by each photo stimulation normalized to the number of photo stimulation trials (in percentage) in tdTomato and ChR2 injected mice (N = 5 and 3 mice, respectively). (**C**) Schematics of photo stimulation of DMH^GAD2^ neurons in the home-like cage with beddings (without TMT). The bar graph shows the number of bedding-pushing occurrences by each photo stimulation normalized to the number of photo stimulation trials (in percentage) in tdTomato and ChR2 injected mice (N = 5 and 4 mice, respectively). (**D**) Example of a double-color in-situ mRNA hybridization in DMH. DAPI is shown in blue and TAC1+ and VGAT+ cells are in green and red, respectively. The bottom right panel shows the overlap between the three colors. Scale bar indicates 0.1 mm. (**E**) Expression of AAV-ChR2-mCherry in DMH of a TAC1-Cre mouse. Scale bar indicates 0.1 mm. Dashed line indicates the fiber position. DMH: Dorsomedial nucleus of hypothalamus, LHA: Lateral hypothalamic area, SubI: subincertus nucleus, VMH: Ventromedial nucleus of hypothalamus, V3: Third ventricle. (**F**) Stacked bar graph shows duration for each action (bedding-pushing, self-grooming, rearing, jumping, tail rattling and freezing) in the home-like cage with beddings (without TMT) in photo stimulated tdTomato and ChR2 TAC1-Cre mice. The bar graph shows the number of bedding-pushing occurrences by each photo stimulation normalized to the number of photo stimulation trials (in percentage) (N = 5 and 12 mice, respectively). (**G**) Z-score (left) and mean Z-score (right) of pupil size of tdTom (red; N =5 mice) and ChR2 (blue; N = 5 mice) mice with photo stimulation from 0 to 5 s. (**H**) Z-score (left) and mean Z-score (right) of whisker activity of mice in (F). (**I**) An example heatmap of the track of a ChR2 TAC1-Cre mouse in a real time place preference/aversion (RTPPA) test. The bar graph shows the duration of time that control tdTom and ChR2 (N = 8 and 8 mice, respectively) mice spent in the opto-linked chamber in the RTPPA test. Red data point indicates the example mouse presented in the heatmap track. (**J**) Schematics of photo stimulation of DMH^TAC1^ neurons in the home-like cage with their own nesting material inside the cage and beddings both inside the cage and outside the cage, on an elevated platform around the cage, to facilitate traveling between inside and outside the cage and to make equal possibility of bedding-pushing behavior in both sides. Mice are photo stimulated all the time (with random time intervals between the pulses), both inside and outside the cage. The bar graph shows the percentage of time spent inside the cage in tdTomato and ChR2 mice (N = 5 and 5, respectively). (**K**) Schematics of the photo stimulation conditioning test, where ChR2 mice on day 2 and day 3 are photo stimulated in the home-like cage. On day 1, pre-conditioning, and day 4, post-conditioning, there is no photo stimulation. The bar graph shows duration of bedding-pushing before and after conditioning (N = 5 mice). (**L**) Schematics of photo inhibition of DMH^TAC1^ neurons in the outside-stressor test with the TMT cottons. The bar graph shows the duration of bedding-pushing in tdTomato and stGtACR2 injected mice (N = 6 and 5 mice, respectively). (**M**) Schematics of photo inhibition of DMH^TAC1^ neurons in the inside-stressor test with the TMT cottons. The bar graph shows the number of jumps in tdTomato and stGtACR2 injected mice (N = 5 and 5 mice, respectively). (**N**) Schematic of calcium photometry of DMH^TAC1^ neurons in inside- and outside-stressor tests. (**O**) Left: Example Calcium activity of DMH^TAC1^ neurons (dF/F0 Z-score) in an outside-stressor test. Blue bar shows duration of push-bedding. Right: Calcium activity of DMH^TAC1^ neurons (dF/F0 Z-score) in inside- (green) and outside-stressor (orange) tests, averaged over trials. The light color around the average indicates the standard error. Time 0 s in inside-stressor test is the onset of running away (or running + jumping) and in outside-stressor test is the onset of bedding-pushing. (**P**) Mean Z-score of calcium activity in response to stressors in outside- and inside-stressor tests (N = 45 and 104 events, 4 mice, respectively). *: p < 0.05, **: p < 0.01, ***: p < 0.001. See also Figure S4, Video S3 and Table S1.

Furthermore, we asked whether activation of the DMH^TAC1^ neurons induces a general state of fear or if it only elicits the bedding-pushing action. To answer this question, we first examined whether a negative arousal level is increased by activation of the DMH^TAC1^ neurons. Therefore, we optogenetically activated DMH^TAC1^ neurons in head-fixed mice and video recorded the pupil and whiskers. The pupil size and whisker activity measures showed an increased arousal level by activation of the DMH^TAC1^ neurons (**Figures 3G and 3H**). The valence was examined using a real-time place preference/avoidance test in a double-chamber, where one of the chambers is linked to the optogenetic light (light-chamber). Compared to the tdTomato control mice, activation of the DMH^TAC1^ neurons significantly decreased the time spent in the light-chamber (**Figure 3I**). These results together showed that activation of the DMH^TAC1^ neurons induces a negative arousal that is associated with fear or stress.

In order to see whether the photo-induced fear is associated with an outside-stressor, we placed the mouse in a cage with bedding and its own nesting material for 10 minutes, while the outside surface is elevated to make it as easy as possible for the mouse to explore outside the cage. Activation of the DMH^TAC1^ neurons during the whole period in this paradigm significantly increased the time spent inside the cage compared to the tdTomato control mice (**Figure 3J**). While the negative valence will be induced by the photo-activation at both locations, the mice thus prefer to do bedding-pushing in the vicinity of their own nesting material and do not try to escape. The behavior induced by photo-activation of DMH^TAC1^ neurons is thus consistent with the behavior in response to an outside-stressor.

Next, we examined whether repetitively activating the DMH^TAC1^ neurons over days can form an associative fear memory of being trapped by the outside-stressor situation. To achieve this, we first put the mice 10 minutes in a cage with bedding and a visually cued context and measured their behavior. For the next two days the DMH^TAC1^ neurons were photostimulated in the cage for 2 minutes. Next day, we put the mice 10 minutes in the cage without photostimulation and measured their behavior again. The results showed a dramatic increase in bedding-pushing (and not other fear responses; i.e. freezing, jumping, tail rattling) in the last day compared to the first day (**Figure 3K**), implying formation of an associative fear memory of being trapped by the outside-stressor situation.

To answer the question whether bedding-pushing behavior is related to aggression, we photostimulated DMH^TAC1^ neurons in these mice in an open field (without bedding) with intruder conspecifics. Mice showed no aggressive behavior (**Figures S4B-S4E**), indicating that the bedding-pushing is not a proactive or defensive aggression. Furthermore, activation of the DMH^TAC1^ neurons decreased the social interactions (**Figures S4B-S4E**). Moreover, further experiments showed that the behavior induced by the photo-activation is not associated with food and object seeking and running to a nest.

Next, we questioned whether the DMH^TAC1^ neurons are necessary for expressing the fear response to an outside-stressor. To answer this, we expressed stGtACR2 in the DMH^TAC1^ neurons and photo-inhibited them in the outside-TMT paradigm. This intensely suppressed the bedding-pushing behavior, compared to tdTomato control mice (**Figure 3L**) and there were no other compensatory stress responses (jumping, freezing, tail-rattling). Furthermore, photo-inhibition of the DMH^TAC1^ neurons did not significantly suppress the number of jumps in the inside-stressor paradigm (**Figure 3M**). Moreover, calcium photometry of GCaMP6s expressed in DMH^TAC1^ neurons (**Figure 3N**) showed that these neurons are activated by stressors in the outside-stressor situation and during bedding-pushing behavior, but not in the inside-stressor situation and not during running/jumping (**Figures 3O and 3P**). Therefore, the DMH^TAC1^ neurons are specialized for the outside-stressor fear and fear response and not for inside-stressor fear and fear response.

### DMH to PAG projections mediate the outside and inside-stressor responses

Periaqueductal gray (PAG) is a key midbrain region in processing the fear response^19,26^. Previous studies have shown that inhibition of PAG neurons or activation of serotonin receptors in PAG reduces the cardiovascular response to emotional stress evoked by the DMH activation^37,38^. Therefore, we examined whether the DMH projection to PAG^39–41^ also coordinates the responses to the inside- and outside-stressors. Interestingly, lateral PAG receives direct input from both DMH^TAC1^ neurons (**Figure 4A**) and DMH^VGluT2^ neurons (**Figure 4B**). Therefore, we asked whether the DMH^TAC1^ → lPAG and DMH^VGluT2^ → lPAG projections contribute to the outside-stressor and inside-stressor fear responses. To answer this, we first expressed ChR2 in DMH^TAC1^ neurons and DMH^VGluT2^ neurons in two groups of mice and optogenetically activated projections of neurons to the lPAG. Photostimulation of the DMH^TAC1^ → lPAG and DMH^VGluT2^

**Figure 4.**
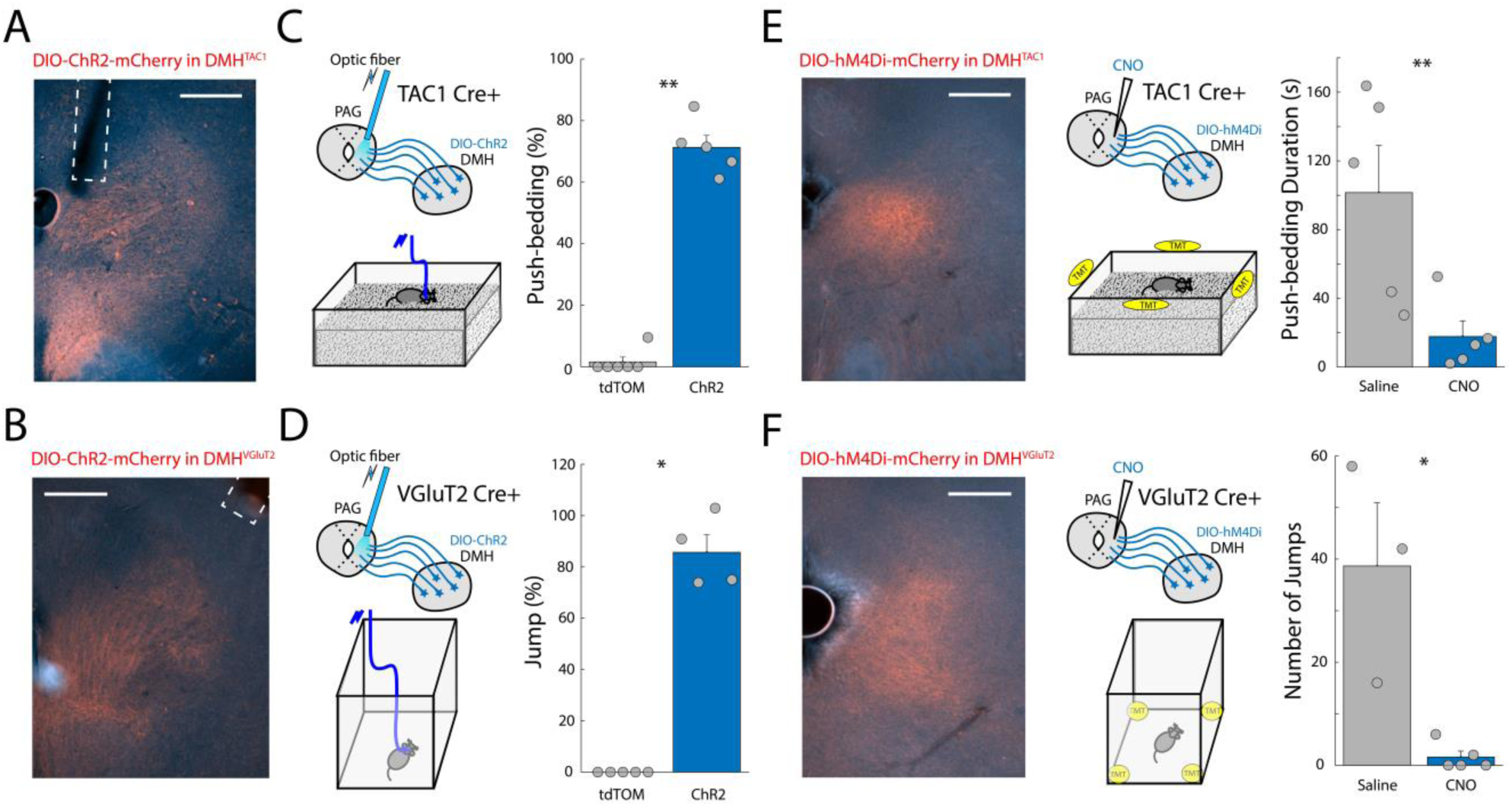
Outside-stressor and inside-stressor responses are controlled through distinct DMH to PAG projections. (**A**) Expression of AAV-ChR2-mCherry in DMH^TAC1^ axons in lPAG. Scale bar indicates 0.25 mm. Dashed line indicates the fiber position. (**B**) Expression of AAV-ChR2-mCherry in DMH^VGluT2^ axons in lPAG. Scale bar indicates 0.25 mm. Dashed line indicates the fiber position. (**C**) Schematics of photo stimulation of DMH^TAC1^ axons in lPAG in the home-like cage with beddings (without TMT). The bar graph shows the number of bedding-pushing occurrences by each photo stimulation normalized to the number of photo stimulation trials (in percentage) in tdTomato and ChR2 injected mice (N = 6 and 5 mice, respectively). (**D**) Schematics of photo stimulation of DMH^VGluT2^ axons in lPAG in the box (without TMT). The bar graph shows the number of jumps by each photo stimulation normalized to the number of photo stimulation trials (in percentage) in tdTomato and ChR2 injected mice (N = 5 and 4 mice, respectively). (**E**) Left: Expression of AAV-hM4Di-mCherry in DMH^TAC1^ axons in lPAG. Middle: Schematics of chemogenetic inhibition of DMH^TAC1^ axons in lPAG in the outside-stressor test with TMT. Right: Bar graph shows the duration of bedding-pushing in mice with local injection of saline and CNO in lPAG (N = 5 and 5 mice, respectively). (**F**) Left: Expression of AAV-hM4Di-mCherry in DMH^VGluT2^ axons in lPAG. Middle: Schematics of chemogenetic inhibition of DMH^VGluT2^ axons in lPAG in the inside-stressor test with TMT. Right: Bar graph shows the number of jumps in mice with local injection of saline and CNO in lPAG (N = 3 and 5 mice, respectively). *: p < 0.05, **: p < 0.01. See also and Table S1.

→ lPAG showed a significant increase in the bedding-pushing (**Figure 4C**) and jumping (**Figure 4D**) behaviors, respectively. Next, we investigated whether these projections are necessary for the outside-stressor and inside-stressor fear responses. In order to test this, using AAV-DIO-hM4Di and local injection of CNO in lPAG, we chemogenetically suppressed the DMH^TAC1^ → lPAG and DMH^VGluT2^ → lPAG axons. Inhibition of DMH^TAC1^ → lPAG axons showed a dramatic reduction of bedding-pushing in the outside-stressor TMT paradigm (**Figure 4E**). Inhibition of DMH^VGluT2^ → lPAG axons showed an intensive decrease in jumping behavior in the inside-stressor TMT paradigm (**Figure 4F**). We conclude that different projections from the DMH to the lPAG drive the response to both inside- and outside-stressors.

## DISCUSSION

We found that in the DMH, a region of hypothalamus previously implicated in other forms of psychological and physical stress responses^15,16,24^, two distinct neuronal populations differentially drive the anxiety associated responses to the outside- and inside-stressors. Projections from DMH^VGluT2^ neurons to lPAG are essential for escape jumping in response to inside-stressors, while projections from DMH^TAC1^ neurons to lPAG are essential for the bedding pushing behavior in response to outside-stressors.

Animal research has well established that defensive responses depend on the proximity of the threat^8,42^. Proximity is not a direct function of distance, but is linked to the intensity and imminence of the threat and is dependent on the presence of escape options. This threat distance is a psycho-spatial cognitive construct that is also applied to human anxiety and fear responses and their pathology^12,43^. While it is clear that the brain’s defensive system responds dynamically based on the proximity of a threat, it is unclear how this psycho-spatial distance is translated into different actions. Different brain areas are supposedly differently activated depending on the proximity^8^. Prefrontal cortex is activated by distant threats and psychological stressors^15,44^. Based on human imaging data, it has been proposed that brain activation is distributed and shifts from prefrontal areas to the midbrain when distal threats become more proximal^43^. This model, however, is not yet supported by finer scale animal data. At the level of the midbrain, in particular the periaqueductal gray, it is clear that distinct parts of the PAG are involved in the different defensive responses that occur for distal or proximal threats^8^. Upstream of the PAG, differential coding of threat distance remains an open question^9^. Here, our novel paradigm with outside-stressor and inside-stressor tests pointed to the DMH as the upstream region with such differential coding. Previous work using lesioning and pharmacological experiments have shown that altering the activity of DMH neurons alters both autonomic and behavioral responses to stressors^45–50^. For instance, autonomic stress responses, including an increase in heart rate, arterial pressure, body temperature, and escape behaviour occur as a result of blocking GABA receptors in the DMH^37,51^. Administration of GABA receptor agonist in the DMH attenuates the cardiovascular stress response^48,50^ and decreases anxiety and inhibits escape performance^22,46^. Blockade of the GABAA receptor lead to coordinated escape jumps^17^. However, all cell types in the DMH were likely to be inhibited by this manipulation. We found that the GABAergic DMH→PAG projection is specifically required for the outside-stressor response, bedding-pushing, and that the glutamatergic DMH→PAG projection is required for the inside- stressor response, escape jumping. The paraventricular nucleus of hypothalamus (PVN) also receives a prominent glutamatergic, GABAergic and, interestingly, substance P input from the DMH^33,52,53^, which could potentially induce autonomic stress responses associated with anxiety disorders. Therefore, in parallel to the DMH→PAG pathway to execute the associated behavioral responses, the DMH→PVN projections may differentially induce autonomic stress responses associated with agoraphobia and claustrophobia. Our findings, however, elaborate the crucial role of the DMH in behavioral flexibility and context-dependent selection of behavioral strategies in response to stressors, beyond merely driving autonomic stress response.

Differentiating these two psycho-spatially distinct contexts, sensing the outside and inside stressors, needs integration of sensory information and a top-down control resulting from the perceptual differentiation. Interestingly, DMH receives a predominant input from the ventromedial prefrontal cortex (vmPFC) ^54,55^ which can exert top-down control over subcortical structures, such as DMH, to regulate anxiety-associated behavioral responses^54^. A recent study showed that an excitatory projection from specific areas of vmPFC to DMH drives autonomic responses to social stress^24^. Therefore, we hypothesize that vmPFC input to DMH sends the perceptually differentiated signals associated with the outside- and inside- stressor situations to select appropriate behavioral responses.

Why do the mice behave the way they do? It is clear why the jump behavior that is selected by the DMH is appropriate. The behavior seems intended on escaping the threatening situation. Its planning is distinct from a startle jump, and when one location offered more chance to exit the arena, the mice would go to that location to make the jump. The nature and goal of the bedding pushing behavior are more mysterious. The shoveling movement of the snout, also called push dig behavior (www.mousebehavior.org) and the vigorous bedding pushing is distinctly different from the light digging that occurs in exploration^44^. It is much more similar to the “swimming” or tunneling through the bedding substrate that occurs for defensive burying of a shock-prod^56^. For this reason, this odorant-induced bedding pushing has previously been labelled defensive burying^57^. Defensive burying, however, is commonly described as goal-directed and specifically directed at burying the shocking prod^21,56^. The TMT-induced bedding pushing behavior appears to lack this goal, and in that sense is more related to the burying behavior seen in the marble burying assay. In this assay, a number of glass marbles gets increasingly buried by bedding shoveled by the animal, but this action is considered to be not directed at burying the marbles^58^ but rather an undirected expression of anxiety-like behavior^44^. The original goal or benefit of the behavior remains to be determined.

Based on our injection location, we studied the anterior part of the dorsomedial hypothalamus^59^. The boundary between the posterior part of the anterior hypothalamus nucleus (AHN) and the anterior end of the DMH, however, is not clearly defined. The transgene expression in the Gad2-cre, VGlut2-cre and TAC1-cre line do not show a clear border between these regions^36^. Without additional markers, it is therefore difficult to pinpoint the circuitry more precisely to the anterior DMH or the posterior AHN. The existing literature does not help to make the distinction with more confidence. Previously, exposure to predators was shown to activate not only DMH^60^, but also AHN^60,61^. DMH was previously implicated in escape behavior^51^, but so was AHN^62^. Escape behavior in general, however, can have many triggers and can also be triggered or modulated by other regions in the hypothalamus, such as the more ventrally located ventromedial hypothalamus (ventral to our injection site)^62^, and the more anteriorly located paraventricular hypothalamic nucleus^63^. Recent electrophysiological findings suggest more generally that anatomically distributed neuron ensembles in the hypothalamus encode fear behavior classes^64^, and thus restricting the location based on escape behavior is unlikely to be possible. The injection location coordinates are therefore perhaps the best indication of the population that we describe in this manuscript.

It is no surprise to find involvement of the PAG in these stress responses. Neurons in the PAG are activated by exposure to predator odor^61,65^ and PAG mediates a range of defensive behaviors^14,66^. The PAG is an extended regio surrounding the mesencephalic aqueduct. The PAG is divided into rostro-caudal columns which have different molecular profiles and behavioral outputs^14,67,68^. The dorsal and dorsolateral part of the PAG (dlPAG) appear to be responsible for initiation of escape behavior^19^, in particular activity of glutamatergic neurons in the dorsal PAG increases selectively during escape initiation^26^. The lateral and dorsolateral PAG have been proposed to be preferentially activated by “escapable” stressors and associated with active defensive responses^66^, while the ventrolateral PAG (vlPAG) is proposed to be activated by “inescapable” stressors for which passive coping behavior is the primary response. This would suggest that input from the DMH^Vglut2^ neurons activates the dlPAG and lPAG to initiate the escape behavior, and while the DMH^GAD2^ neurons perhaps inhibit the escape behavior and a disinhibitory circuits leads to the bedding pushing response. A disinhibitory circuit motif has been shown in the connectivity of the central nucleus of the amygdala (CeA) to the vlPAG, where inhibition from the CeA leads to initiation of a freezing response^69^. Along the rostro-caudal axis, there is also heterogeneity in the PAG in connectivity and molecular expression^67,68^. Our experimental manipulations were aimed at the center of the PAG along this axis, but the exact position along this axis can have an impact on behavior which remains to be determined^68,70^.

In summary, we showed that in the DMH two cell-types are differently activated by two different psycho-spatial distances of the threat, and the presence of an escape possibility. These two situations are the source of extreme anxiety in agoraphobia and claustrophobia and the evolutionary conservation of the mammalian defensive system suggests that in these disorders these different populations of DMH neurons will be activated. This gives a new handle to investigate these disorders and test possible therapies.

## ACKNOWLEDGEMENTS

We thank Mejdy Kacem, Andres de Groot, Ralph Hamelink, Maurice Heemskerk, Martin Oomstee, Stephen Super, John van Veldhuijzen, Marian Verhage, Nanneke van der Wal, Joris Coppens and Joost Verhaagen. **Funding:** This work was supported by the Dr. J.L. Dobberke Foundation. **Competing interests:** Authors declare no competing interests. **Data and materials availability:** All behavioral videos and scripts are stored on institutional storage and are available upon request from the authors.

## AUTHOR CONTRIBUTIONS

M.A. conceived the study with input from J.A.H. M.A., M.G., J.F.M.v.V., T.v.L., W.B., J.H.W.H., M.Y.S., J.L.C., M.D., M.H.P.K., R.H. and J.A.H. collected and analyzed the data. F.d.W. made viral vectors and helped the in-situ hybridization. M.A. and J.A.H. wrote the paper.

## STAR METHODS

### RESOURCE AVAILABILITY

#### Lead contact

Further information and requests for resources and reagents should be directed to and will be fulfilled by the lead contact Alexander Heimel.

#### Materials availability

This study did not generate new unique reagents.

#### Data and code availability

All data reported in this paper will be shared by the lead contacts upon request. Custom-written codes are available on GitHub repository https://github.com/heimel. Any additional information required to reanalyze the data reported in this paper is available from the lead contacts upon request.

### EXPERIMENTAL MODEL AND SUBJECT DETAILS

#### Mice

Mice were housed under controlled climate (22-24°C) in a normal light/dark cycle (12 hr / 12 hr) with ad libitum access to laboratory food pellets and water. C57BL/6J (Janvier) mice, Gad2-Cre (Stock #028867, Jackson), VGlut2-Cre (Stock # 016963, Jackson), and TAC1-Cre (Stock #021877, Jackson) mice of 2–5 months of age from either sex were used for the experiments. All experimental protocols were approved by the institutional animal care and use committee of the Royal Netherlands Academy of Sciences (KNAW) and were in accordance with the Dutch Law on Animal Experimentation. Reporting has followed ARRIVE guidelines.

### METHOD DETAILS

#### Virus vector injection

Mice were anesthetized with isoflurane (5% induction, 1.2–1.8% maintenance) in oxygen (0.6 L per min flow rate). Body temperature was maintained at 36.5 ℃, using a controlled heating pad (Harvard Apparatus). The eyes were protected from light by black stickers and from drying by Cavasan eye ointment. During the surgery, mice were administered the analgesic Metacam (1 mg per kg s.c.) to reduce pain during the recovery. Using ear bars, mice were head fixed on a stereotactic device (Kopf) and using a scalpel blade the scalp along with the midline was cut to expose the skull. Small craniotomies were made by dental drill and using a Drummond Nanoject volume injector the virus (80 nl) was injected into the target brain regions. Fifteen minutes after the injection, the glass pipette was retracted and in case we did not need fiber implantation, the scalp was sutured. After recovery, the animals were returned to their cage.

Brain regions and coordinates (from Bregma) used for virus injections: anterior DMH (AP: -1.4 mm, ML: 0.3 mm, DV: 4.9 mm) and PAG (AP: -4.5 mm, ML: 0.5 mm, DV: 2.2-2.6 mm). For optogenetic experiments, we used AAV9-hEF1a-DIO-mCherry-hChR2 (University of Zurich; V80-9, a gift from Karl Deisseroth), AAV1-hSyn-SIO-FusionRed-stGtACR2 ^71^ (Addgene viral prep #105677-AAV1, a gift from Ofer Yizhar), for chemogenetic experiments, AAV1-hSyn-DIO-mCherry-hM4Di (made in the host institute), and for fiber photometry experiments, AAV1-Syn-FLEX-GCaMP6s^72^ (University of Pennsylvania, a gift from Douglas Kim & GENIE Project).

#### Optic fiber implantation

For optogenetic behavior tests and fiber photometry, mice were fiber and head plate implanted during the virus injection surgery. After the virus injection, the scalp and soft tissue overlying the skull were incised to expose the skull. Craniotomies with 300-400 µm diameter were made (by dental drill) to insert the optic fibers (200 µm diameter). After cementing a metal ring (9 mm inner diameter) to the skull, the optic fibers were inserted 300 µm (for optogenetics; fibers with N.A. of 0.22) or 50 µm (for fiber photometry; with N.A. of 0.57) above the target brain regions and cemented to the skull. After recovery, the animals were returned to their cage. Positions and coordinates (from Bregma) used for optic fiber implantation: anterior DMH (AP: -1.4 mm, ML: 0.9 mm, ML-angle: 7°, DV for photometry: 4.85 mm, DV for optogenetics: 4.60 mm) and PAG (AP: -4.5 mm, ML: 1.0 mm, ML-angle: 14°, DV for optogenetics: 2.0-2.1 mm).

#### Behavior tests and analysis

Single-housed mice were habituated to the experimenters for three days (30 min a day). Additionally, one day (for 15 min) and 10 min prior to the experiment, mice were habituated to the experimental box (20 cm × 20 cm × 30 cm (h)) for the inside-stressor experiments and to the cage (23 × 17 × 14 cm, L x W x H) with bedding (corn cob GM12) for outside-stressor experiments. For the inside-stressor test, mice were placed in the box with cottons contaminated with 5 µl TMT (BioSRQ: purity > 90.0%) secured in all corners (**Figure 1A**). For the outside-stressor test, mice were put in an open cage (identical to their own home-cage) with bedding (9 cm height) with cottons contaminated with 5 µl (or 3 µl where it is mentioned in the text) TMT secured around and outside the cage (**Figure 1D**). The behavior was video recorded for 10 min. Each mouse was exposed to the TMT only in one experimental session (e.g. in **Figure 1C** each condition represents a separate group of mice).

Videos of the trials were manually labeled frame by frame using JAABA^73^. Analyzers were blind to the experimental groups of the mice. The behaviors displayed during inside-stressor test were categorized as: jumping, running (running-away upon approaching the stressors), self-grooming (grooming whiskers, face, body, or tail), rearing (upward), tail-rattling and freezing (startle immobility). In photo activation experiments, additional categories were included as well: alertness (stopping the continuing behavior and orienting the head), running, posture (making jumping posture; stretching the body with pressure on the limbs while the head is upward). The behaviors displayed during outside-stressor test were categorized as: push-bedding (pushing forward the bedding materials while the snout is in the beddings, mainly by snout and partially by forepaws (**Video S1**), self-grooming, rearing (upward), and stereotypic defensive responses, i.e. jumping, tail-rattling and freezing (startle immobility). The JAABA labeling was imported to MATLAB for further analysis (using custom-written MATLAB programs).

#### Optogenetics and chemogenetics

For induction of fear response to inside- and outside-stressor, the bilateral photo activation was done by laser (465 nm; 3 mW; Shanghai Laser & Optics Century Co.) pulses switching at 20 Hz with varied pulse duration (1-4 s) and at random time points, unless it is mentioned otherwise in the text. Photo inhibition was done by laser (465 nm; 3 mW) shining continuously (0 Hz) throughout the test duration in presence of inside- or outside-stressors. In case we needed to bilaterally silence hM4Di-expressed axon terminals, we locally injected 100 nl of CNO-dihydrochloride (10 µM) in the target region (where the craniotomy was made above the PAG during the virus injection + head plate implantation surgery) 30 minutes before the test, using a Drummond Nanoject volume injector, while the animals where briefly head-fixed. All the optogenetic and chemogenetic manipulations were done bilaterally. Up to a few weeks after the behavioral tests, mice were perfused with 4% PFA in PBS and brains were dissected for imaging and histology.

#### Calcium fiber photometry and data analysis

Mice were habituated to the experimenters for three days (30 min a day) and the experimental box (35 cm × 35 cm × 35 cm) for one day (15 min). At least 3 weeks after the GCaMP6 viral injection and the optic fiber implantation, calcium activity of DMH neurons was recorded by a fiber photometry system (Doric Lenses) in freely moving mice in the outside- and inside-stressor paradigms. The behavior videos and calcium transients were recorded simultaneously using Raspberry Pi cameras (frame rate: 30 fps) and the fiber photometry system (PyPhotometry; sampling rate: 100 Hz), respectively. The excitation blue light (LED; 465 nm) at the tip of the optic fiber was adjusted around 30 µW. The data for fiber photometry was analyzed using a custom-written MATLAB program. Behavior of the mouse and stress responses were analyzed using JAABA, similarly to the optogenetic experiments. Onset of the behavior labels was considered as time zero to calculate the values of calcium transients change (Z-score): onset of orienting/running away for inside-stressor paradigm and onset of push-bedding for outside-stressor paradigm. A linear drift correction was applied. The mean baseline was taken as the average signal of the entire recording session. Z-score was calculated by subtracting the mean baseline and further dividing by the standard deviation of the baseline distribution. After the photometry experiments, mice were perfused with 4% PFA in PBS and brains were dissected for imaging and histology.

#### Pupil size, whisker activity and data analysis

For these experiments, mice were habituated for three days (1 hr a day) to the head-fixed setup. At least 3 weeks after the virus injection/fiber implantation surgery, mice with expression of tdTomato (control) or ChR2 in the DMH were used for measuring the change in physiological arousal level caused by the photostimulation in a room with dim ambient light. Mice were head-fixed using a magnet and an infrared LED light was directed to the face (mounted 10 cm away). The implanted fibers were coupled to a 465 nm laser (3 mW). Mice received 20 Hz photostimulation with 5 s pulses (20 pulses with 15 s intervals). We monitored the effect of photostimulation on the arousal level by recording pupil and whisker videos (IR camera: Basler acA640-90um; with frame rate of 25 fps). Whisker activity of each frame was calculated by absolute value of subtraction of the averaged intensity of that frame from the previous frame. The pupil size and whisker activity data were measured using a custom-written MATLAB program.

#### Real-time place preference/avoidance test (RTPPT)

At least 3 weeks after the virus injection/fiber implantation surgery, mice with expression of ChR2 or tdTomato (control) in DMH^TAC1^ or DMH^VGlut2^ were used for real-time place preference/avoidance test in a two-chamber white box (60 cm × 30 cm × 30 cm) without additional contextual cues. After 10 min habituation to the box, one chamber was paired with a 20 Hz photostimulation and the other identical chamber had no photostimulation. The laser-coupled chamber was randomly assigned. The total test duration was 10 min. Mice tracks were analyzed using Bonsai software (https://open-ephys.org/bonsai).

#### Opto-conditioning RTPPT test

Mice with expression of ChR2 or tdTomato (control) in DMH^VGlut2^ were used for testing associative fear memory in an RTPPT double-chamber box, where the chambers were visually distinct (by having different patterns on the walls). On the first day, mice were in the box for 10 min without photostimulation. On the second and third days, mice received 20 Hz photostimulation in their preferred chamber. The next day, again mice were in the box for 10 min without photostimulation. The fraction of time mice spent in the chamber that was not preferred on the first day was measured and compared to having no preference.

#### Opto-conditioning test in home-like cage

Mice with expression of ChR2 or tdTomato (control) in DMH^TAC1^ were used for testing associative outside-stressor memory in a home-like cage with a thick layer of beddings, where the outside walls were taped black to make it more distinct from the home-cage. On the first day, mice were in the home-like cage for 10 min without photostimulation. On the second and third days, mice received 20 Hz photostimulation in the home-like cage. The next day, again mice were in the home-like cage for 10 min without photostimulation. The duration of push-bedding on the first and last days was measured and compared.

#### Social interaction test

Single-housed mice were habituated to the experimenters and the experimental box (35 cm × 35 cm × 35 cm) every day for three weeks. A novel conspecific (‘intruder’) (with the same sex and weight (± 1 grams)) was placed in the box after the ‘resident’ mouse had been there for 30 min. The intruder was always a new mouse for the ‘resident’ mouse that had been in the box. The behavior was video recorded for 10 min. Photo activation was done by laser (465 nm; 3 mW at the tip of the optic fiber; Shanghai Laser & Optics Century Co.) switching at 20 Hz with 50% duty cycle throughout the test duration.

Videos of the trials were manually labeled frame by frame using JAABA. Analyzers were blind to the experimental groups of mice. The behaviors displayed during social interaction were categorized as: Acceptance: when the resident mouse accepts to stay (without taking any investigatory interaction), while it is investigated by the intruder mouse. Active interaction: when the resident mouse actively approaches the intruder mouse initiates the interaction and investigates the intruder mouse. Passive interaction: when the interaction of the resident mouse with the intruder mouse starts with approach and initiation of the intruder mouse. Interaction includes body, facial, and anogenital touching/sniffing and grabbing/holding. Aversion: when the resident mouse shows aversive behavior in response to the intruder mouse, such as running away and tail rattling. Aggression: when the resident mouse shows aggressive behaviors such as fighting, boxing, wrestling and biting. The JAABA labeling was imported to MATLAB for further analysis (using custom-written MATLAB programs).

#### CFos staining

For cFos staining, 4 groups of each 4 mice were used. All groups were habituated to the experimenter as described for the other behavior tests. The first group underwent the inside-stressor paradigm in a box with TMT-cottons attached to all four corners inside the cage. The second group underwent the same procedure, but without TMT on the cottons, as a control group. The third group underwent the outside-stressor paradigm in a cage with a high level of bedding with TMT-cottons attached around the outside of the cage. The fourth group underwent the same procedure, but without TMT on the cottons, as a control. Sixty minutes after the test, mice were taken out of the experimental box and immediately were transcardially perfused with 4% paraformaldehyde (PFA) in phosphate buffered saline (PBS). The brains were extracted and post-fixated for 2 hours in PFA at 4 °C before moving to a PBS solution. Later, we cut the middle part of the brain (including hypothalamus and thalamus) in coronal slices of 50 μm thickness on a vibratome and stained the slices with a cFos (9F6) Rabbit mAb primary antibody (Cell Signaling Technology, 1:2500) and a goat-antirabbit Alexa 488 secondary antibody (Invitrogen, 1:500). Next, all slices were mounted using mowiol containing DAPI and imaged on a Zeiss Axio Scan Z1 slide scanner at 10x. The segmentation toolkit Ilastik^74^ was trained on a random set of image parts per mouse and used to count cFos positive neurons across the entire slices. All slices and cells detected by Ilastik were aligned to the Allen Common Coordinate Framework by Amasine^75^. Finally, the results of all hemispheres from each group were combined and the imaged brain was divided in cubic voxels of 0.25 mm width. No distinction was made between the left and right hemisphere. For each voxel, t-tests over all hemispheres were performed to compare inside-TMT condition with the no-TMT condition, and the outside-TMT condition with its control.

#### Multi-fluorescence mRNA in-situ hybridization (RNAscope)

In-situ hybridization was performed using RNAscope Technology. We used C57BL/6 mice for these experiments. After induction of deep-anesthesia by isoflurane (5%), brains were extracted and immediately fresh-frozen in optimal cutting temperature (OCT). Using a cryostat, brains were sliced into 10 µm sections, mounted on glass slides, and stored at -80 °C. Multi-fluorescence mRNA situ hybridization was performed using ACDBio RNAscope multiplex fluorescence assay (https://acdbio.com/). The RNAscope protocol was carried out as indicated in the user manual of the ACDBio RNAscope. The brain sections of DMH were post-fixed with 4% chilled paraformaldehyde (PFA in PBS) for 15 min and then dehydrated through four dehydration steps in 50%, 75%, 100%, and 100% ethanol (5-min each), respectively, at room temperature (RT). After air drying for 5 min at RT, Protease IV was applied to the slices for 35 min at RT. Then, they were 3 times washed out by rinsing in phosphate-buffered saline (PBS). TAC1-C1 (Cat No. 410351) and VGAT-C2 (Cat No. 319191-C2) were pipetted onto each slice. Probe hybridization took place in an oven set to 40 °C for 2 hr, and then, slices were rinsed in 1 × RNAscope wash buffer. After incubation at 40 °C in four-step amplifications (Amp1 for 30 min, Amp2 for 15 min, Amp3 for 30 min, and Amp4 for 15 min), slices were mounted with DAPI (4’,6-diamidino-2-phenylindole) vectashield (Vector Laboratories). Immediately after mounting, the stained slices were imaged by confocal SP8 microscope (Leica) using a 20X objective.

### QUANTIFICATION AND STATISTICAL ANALYSIS

#### Overall experimental design and analysis

No statistical methods were used to predetermine sample sizes, but our sample sizes were determined based on previous studies^24,76^. The order of the animals in different experimental groups was randomly assigned. Experimenters were not blind to the experimental conditions, but the collected data were encoded blindly (using a MATLAB code) and analyzers were blind to the experimental conditions. All data were analyzed using JAABA, MATLAB and Bonsai.

#### Statistical analysis

Data are represented as mean ± SEM. All the statistical analysis was done using a homemade MATLAB toolbox, InVivoTools^77–79^ (https://github.com/heimel/InVivoTools). First, normality of the data distribution was checked, using Shapiro-Wilk normality test. In order to assess group statistical significance, in case the data were normally distributed we used parametric tests (i.e. t-test and paired t-test for non-paired and paired comparisons, respectively) and otherwise non-parametric tests (i.e. Mann-Whitney U test and Wilcoxon signed rank test for non-paired and paired comparisons, respectively), followed by a Bonferroni p-value correction for multi group comparisons. To compare the photometry signals in Figures 2 and 3 we used two-way ANOVA followed by a Tukey-Kramer post hoc test. To test for time-locking of the calcium signal to behavioral events we used the time-series zeta test^80^. Individual data points are shown in the figures. All the statistics used in figures and supplementary figures are shown in table of statistics (**Table S1**).

## Supplemental table/video legends

**Table S1. Statistics of main and supplementary figures. Related to Figures 1-4**.

**Video S1. A wild-type mouse in inside-stressor and outside-stressor tests. Related to Figure 1**.

**(A)** Example of behavior of a wild-type mouse in the inside-stressor test with TMT in all corners. The color of the square indicates the behavior during that frame. Blue: jumping. (**B**) Example of behavior of a wild-type mouse in the inside-stressor test with TMT in two corners. (**C**) Example of behavior of a wild-type mouse in the outside-stressor test with TMT. The color of the square indicates the behavior during that frame. Blue: push-bedding; yellow: rearing. (**D**) Example of behavior of a wild-type mouse in the outside-stressor test with air puff.

**Video S2. Activation of DMH^VGluT2^ neurons. Related to Figure 2**.

**(A)** Example of a mouse behavior during photo-activation of DMH^VGluT2^ neurons in an open field without bedding. (**B**) Example of a mouse behavior during photo-activation of DMH^VGluT2^ neurons in an open field with bedding. (**C**) Example of a mouse behavior during photo-activation of DMH^VGluT2^ neurons in a cage with bedding. (**D**) Example of a mouse behavior during photo-activation of DMH^VGluT2^ neurons with different durations of photo-stimulation. The color of the square indicates the behavior during that frame. Yellow: alertness; green: running; light blue: preparatory posture for jumping; dark blue: jumping.

**Video S3. Activation of DMH^TAC1^ neurons. Related to Figure 3**.

Example of a mouse behavior during photo-activation of DMH^TAC1^ neurons in a cage with bedding.

## REFERENCES

1. Masel, C. N., Terry, D. J. & Gribble, M. (1996). The effects of coping on adjustment: Re-examining the goodness of fit model of coping effectiveness. Anxiety, Stress Coping 9.

2. Löw, A., Weymar, M. & Hamm, A. O. (2015). When Threat Is Near, Get Out of Here: Dynamics of Defensive Behavior During Freezing and Active Avoidance. Psychol. Sci. 26.

3. Perna, G., Daccò, S., Menotti, R. & Caldirola, D. (2011). Antianxiety medications for the treatment of complex agoraphobia: Pharmacological interventions for a behavioral condition. Neuropsychiatric Disease and Treatment vol. 7 at 10.2147/ndt.s12979.

4. Wittchen, H. U., Gloster, A. T., Beesdo-Baum, K., Fava, G. A. & Craske, M. G. (2010). Agoraphobia: A review of the diagnostic classificatory position and criteria. Depression and Anxiety vol. 27 at 10.1002/da.20646.

5. Lourenco, S. F., Longo, M. R. & Pathman, T. (2011). Near space and its relation to claustrophobic fear. Cognition 119.

6. Moradi, D., Rahimi, F., Eyvazpour, R., Jahan, A., Rasta, S.H. & Esmaeili, M. (2022). Neural activity in self-identified claustrophobic individuals under in-vivo stimuli: A human electroencephalography dataset. Data Br. 40.

7. Shin, L. M. & Liberzon, I. (2010). The neurocircuitry of fear, stress, and anxiety disorders. Neuropsychopharmacology vol. 35 at 10.1038/npp.2009.83.

8. Fanselow, M. S. (1994). Neural organization of the defensive behavior system responsible for fear. Psychon. Bull. Rev. 1.

9. Mobbs, D., Headley, D. B., Ding, W. & Dayan, P. (2020). Space, Time, and Fear: Survival Computations along Defensive Circuits. Trends in Cognitive Sciences vol. 24 at 10.1016/j.tics.2019.12.016.

10. Stein, M. D. & Steckler, T. (2012). Behavioural Neurobiology of Anxiety and Its Treatment. (Springer).

11. Gross, C. T. & Canteras, N. S. (2012). The many paths to fear. Nature Reviews Neuroscience vol. 13 at 10.1038/nrn3301.

12. McNaughton, N. & Corr, P. J. (2004). A two-dimensional neuropsychology of defense: Fear/anxiety and defensive distance. Neuroscience and Biobehavioral Reviews vol. 28 at 10.1016/j.neubiorev.2004.03.005.

13. Canteras, N. S. (2002). The medial hypothalamic defensive system: Hodological organization and functional implications. Pharmacol. Biochem. Behav. 71, 481–491.

14. Silva, B. A., Gross, C. T. & Gräff, J. (2016). The neural circuits of innate fear: Detection, integration, action, and memorization. Learning and Memory vol. 23 at 10.1101/lm.042812.116.

15. Nakamura, K., Nakamura, Y. & Kataoka, N. (2022). A hypothalamomedullary network for physiological responses to environmental stresses. Nature Reviews Neuroscience vol. 23 at 10.1038/s41583-021-00532-x.

16. Canteras, N. S., Chiavegatto, S., Ribeiro Do Valle, L. E. & Swanson, L. W. (1997). Severe reduction of rat defensive behavior to a predator by discrete hypothalamic chemical lesions. Brain Res. Bull. 44.

17. Brandao, M. L., Di Scala, G., Bouchet, M. J. & Schmitt, P. (1986). Escape behavior produced by the blockade of glutamic acid decarboxylase (GAD) in mesencephalic central gray or medial hypothalamus. Pharmacol. Biochem. Behav. 24.

18. Silva, B.A., Mattucci, C., Krzywkowski, P., Murana, E., Illarionova, A., Grinevich, V., Canteras, N.S., Ragozzino, D. & Gross, C.T. (2013). Independent hypothalamic circuits for social and predator fear. Nat. Neurosci. 16.

19. Lefler, Y., Campagner, D. & Branco, T. (2020). The role of the periaqueductal gray in escape behavior. Current Opinion in Neurobiology vol. 60 at 10.1016/j.conb.2019.11.014.

20. Rosen, J. B., Asok, A. & Chakraborty, T. (2015). The smell of fear: Innate threat of 2,5-dihydro-2,4,5-trimethylthiazoline, a single molecule component of a predator odor. Frontiers in Neuroscience vol. 9 at 10.3389/fnins.2015.00292.

21. Boer, S. F. De & Koolhaas, J. M. (2003). Defensive burying in rodents: ethology, neurobiology and psychopharmacology. Eur. J. Pharmacol. 463, 145–161.

22. Nascimento, J. O. G., Zangrossi, H. & Viana, M. B. (2010). Effects of reversible inactivation of the dorsomedial hypothalamus on panic-and anxiety-related responses in rats. Brazilian J. Med. Biol. Res. 43.

23. Ullah, F., dos Anjos-Garcia, T., dos Santos, I. R., Biagioni, A. F. & Coimbra, N. C. (2015). Relevance of dorsomedial hypothalamus, dorsomedial division of the ventromedial hypothalamus and the dorsal periaqueductal gray matter in the organization of freezing or oriented and non-oriented escape emotional behaviors. Behav. Brain Res. 293, 143–152.

24. Kataoka, N., Shima, Y., Nakajima, K. & Nakamura, K. (2020). A central master driver of psychosocial stress responses in the rat. Science (80-.). 367, 1105–1112.

25. Biagioni, A.F., dos Anjos-Garcia, T., Ullah, F., Fisher, I.R., Falconi-Sobrinho, L.L., de Freitas, R.L., Felippotti, T.T. & Coimbra, N.C. (2016). Neuroethological validation of an experimental apparatus to evaluate oriented and non-oriented escape behaviours: Comparison between the polygonal arena with a burrow and the circular enclosure of an open-field test. Behav. Brain Res. 298, 65–77.

26. Evans, D.A., Stempel, A.V., Vale, R., Ruehle, S., Lefler, Y. & Branco, T. (2018). A synaptic threshold mechanism for computing escape decisions. bioRxiv 272492 doi:10.1101/272492.

27. Steuernagel, L., Lam, B.Y., Klemm, P., Dowsett, G.K., Bauder, C.A., Tadross, J.A., Hitschfeld, T.S., del Rio Martin, A., Chen, W., De Solis, A.J. et al. (2022). HypoMap—a unified single-cell gene expression atlas of the murine hypothalamus. Nat. Metab. 4, 1402-1419.

28. Stotz-Potter, E. H., Willis, L. R. & DiMicco, J. A. (1996). Muscimol acts in dorsomedial but not paraventricular hypothalamic nucleus to suppress cardiovascular effects of stress. J. Neurosci. 16, 1173–1179.

29. De Matteo, R., Head, G. A. & Mayorov, D. N. (2006). Tempol in the dorsomedial hypothalamus attenuates the hypertensive response to stress in rabbits. Am. J. Hypertens. 19, 396–402.

30. Ebner, K. & Singewald, N. (2006). The role of substance P in stress and anxiety responses. in Amino Acids vol. 31.

31. Iftikhar, K., Siddiq, A., Baig, S. G. & Zehra, S. (2020). Substance P: A neuropeptide involved in the psychopathology of anxiety disorders. Neuropeptides vol. 79 at 10.1016/j.npep.2019.101993.

32. Lein, E.S., Hawrylycz, M.J., Ao, N., Ayres, M., Bensinger, A., Bernard, A., Boe, A.F., Boguski, M.S., Brockway, K.S., Byrnes, E.J. et al. (2007). Genome-wide atlas of gene expression in the adult mouse brain. Nature 445.

33. Womack, M. D. & Barrett-Jolley, R. (2007). Activation of paraventricular nucleus neurones by the dorsomedial hypothalamus via a tachykinin pathway in rats. Exp. Physiol. 92, 671–676.

34. Li, Y., Lopez-Huerta, V.G., Adiconis, X., Levandowski, K., Choi, S., Simmons, S.K., Arias-Garcia, M.A., Guo, B., Yao, A.Y., Blosser, T.R. et al. (2020). Distinct subnetworks of the thalamic reticular nucleus. Nature 583.

35. Ahmadlou, M., Houba, J.H., van Vierbergen, J.F., Giannouli, M., Gimenez, G.A., van Weeghel, C., Darbanfouladi, M., Shirazi, M.Y., Dziubek, J., Kacem, M., et al. (2021). A cell type–specific cortico-subcortical brain circuit for investigatory and novelty-seeking behavior. Science 372(6543), eabe9681.

36. Oh, S.W., Harris, J.A., Ng, L., Winslow, B., Cain, N., Mihalas, S., Wang, Q., Lau, C., Kuan, L., Henry, A.M. et al. (2014). A mesoscale connectome of the mouse brain. Nature 508.

37. Da Silva, L. G., Alvim De Menezes, R. C., Souza dos Santos, R. A., Campagnole-Santos, M. J. & Peliky Fontes, M. A. (2003). Role of periaqueductal gray on the cardiovascular response evoked by disinhibition of the dorsomedial hypothalamus. Brain Res. 984, 206–214.

38. Villela, D. C., da Silva Junior, L. G. & Fontes, M. A. P. (2009). Activation of 5-HT receptors in the periaqueductal gray attenuates the tachycardia evoked from dorsomedial hypothalamus. Auton. Neurosci. Basic Clin. 148.

39. Horst, G. J. T. & Luiten, P. G. M. (1986). The projections of the dorsomedial hypothalamic nucleus in the rat. Brain Res. Bull. 16.

40. Thompson, R. H., Canteras, N. S. & Swanson, L. W. (1996). Organization of projections from the dorsomedial nucleus of the hypothalamus: A PHA-L study in the rat. J. Comp. Neurol. 376.

41. de Git, K.C., van Tuijl, D.C., Luijendijk, M.C., Wolterink-Donselaar, I.G., Ghanem, A., Conzelmann, K.K. & Adan, R.A. (2018). Anatomical projections of the dorsomedial hypothalamus to the periaqueductal grey and their role in thermoregulation: a cautionary note. Physiol. Rep. 6, e13807.

42. Blanchard, D. C. & Blanchard, R. J. (1990). Behavioral correlates of chronic dominance-subordination relationships of male rats in a seminatural situation. Neurosci. Biobehav. Rev. 14.

43. Hamm, A. O. (2020). Fear, anxiety, and their disorders from the perspective of psychophysiology. Psychophysiology 57.

44. Lustberg, D.J., Liu, J.Q., Iannitelli, A.F., Vanderhoof, S.O., Liles, L.C., McCann, K.E. & Weinshenker, D. (2022). Norepinephrine and dopamine contribute to distinct repetitive behaviors induced by novel odorant stress in male and female mice. Horm. Behav. 144.

45. Soltis, R. P. & DiMicco, J. A. (1991). GABA(A) and excitatory amino acid receptors in dorsomedial hypothalamus and heart rate in rats. Am. J. Physiol. - Regul. Integr. Comp. Physiol. 260.

46. Shekhar, A. (1993). GABA receptors in the region of the dorsomedial hypothalamus of rats regulate anxiety in the elevated plus-maze test. I. Behavioral measures. Brain Res. 627, 9–16.

47. Inglefield, J. R., Schwarzkopf, S. B. & Kellogg, C. K. (1994). Alterations in behavioral responses to stressors following excitotoxin lesions of dorsomedial hypothalamic regions. Brain Res. 633, 151–161.

48. Morin, S. M., Stotz-Potter, E. H. & Dimicco, J. A. (2001). Injection of muscimol in dorsomedial hypothalamus and stress-induced Fos expression in paraventricular nucleus. Am. J. Physiol. - Regul. Integr. Comp. Physiol. 280, 1276–1284.

49. Gasser, P. J., Lowry, C. A. & Orchinik, M. (2006). Corticosterone-sensitive monoamine transport in the rat dorsomedial hypothalamus: Potential role for organic cation transporter 3 in stress-induced modulation of monoaminergic neurotransmission. J. Neurosci. 26, 8758–8766.

50. Hunt, J. L., Zaretsky, D. V., Sarkar, S. & DiMicco, J. A. (2010). Dorsomedial hypothalamus mediates autonomic, neuroendocrine, and locomotor responses evoked from the medial preoptic area. Am. J. Physiol. - Regul. Integr. Comp. Physiol. 298, 130–140.

51. Freitas, R. L., Uribe-Mariño, A., Castiblanco-Urbina, M. A., Elias-Filho, D. H. & Coimbra, N. C. (2009). GABAA receptor blockade in dorsomedial and ventromedial nuclei of the hypothalamus evokes panic-like elaborated defensive behaviour followed by innate fear-induced antinociception. Brain Res. 1305, 118–131.

52. Cullinan, W. E., Ziegler, D. R. & Herman, J. P. (2008). Functional role of local GABAergic influences on the HPA axis. Brain Structure and Function vol. 213 at 10.1007/s00429-008-0192-2.

53. Ulrich-Lai, Y. M., Jones, K. R., Ziegler, D. R., Cullinan, W. E. & Herman, J. P. (2011). Forebrain origins of glutamatergic innervation to the rat paraventricular nucleus of the hypothalamus: Differential inputs to the anterior versus posterior subregions. J. Comp. Neurol. 519.

54. McDougall, S. J., Widdop, R. E. & Lawrence, A. J. (2004). Medial prefrontal cortical integration of psychological stress in rats. Eur. J. Neurosci. 20.

55. Giustino, T. F. & Maren, S. (2015). The role of the medial prefrontal cortex in the conditioning and extinction of fear. Frontiers in Behavioral Neuroscience vol. 9 at 10.3389/fnbeh.2015.00298.

56. Pinel, J. P. & Treit, D. (1978). Burying as a defensive response in rats. J. Comp. Physiol. Psychol. 92.

57. Hwa, L.S., Neira, S., Pina, M.M., Pati, D., Calloway, R. & Kash, T.L. (2019). Predator odor increases avoidance and glutamatergic synaptic transmission in the prelimbic cortex via corticotropin-releasing factor receptor 1 signaling. Neuropsychopharmacology 44.

58. Gyertyan, I. (1995). Analysis of the marble burying response: Marbles serve to measure digging rather than evoke burying. Behav. Pharmacol. 6.

59. Franklin, K. B. J. & Paxinos, G. (2007). The Mouse Brain in Stereotaxic Coordinates (map). (Academic Press).

60. Dielenberg, R. A. & McGregor, I. S. (2001). Defensive behavior in rats towards predatory odors: A review. Neuroscience and Biobehavioral Reviews vol. 25 at 10.1016/S0149-7634(01)00044-6.

61. Cezario, A. F., Ribeiro-Barbosa, E. R., Baldo, M. V. C. & Canteras, N. S. (2008). Hypothalamic sites responding to predator threats - The role of the dorsal premammillary nucleus in unconditioned and conditioned antipredatory defensive behavior. Eur. J. Neurosci. 28.

62. Wang, L., Chen, I. Z. & Lin, D. (2015). Collateral Pathways from the Ventromedial Hypothalamus Mediate Defensive Behaviors. Neuron 85.

63. Mangieri, L.R., Jiang, Z., Lu, Y., Xu, Y., Cassidy, R.M., Justice, N., Xu, Y., Arenkiel, B.R. & Tong, Q. (2019). Defensive behaviors driven by a hypothalamic-ventral midbrain circuit. eNeuro 6.

64. Stagkourakis, S., Spigolon, G., Marks, M., Feyder, M., Kim, J., Perona, P., Pachitariu, M. & Anderson, D.J. (2023). Distributed representations of innate behaviors in the hypothalamus do not predict specialized functional centers. bioRxiv 1–65.

65. Canteras, N. S. & Goto, M. (1999). Fos-like immunoreactivity in the periaqueductal gray of rats exposed to a natural predator. Neuroreport 10.

66. Keay, K. A. & Bandler, R. (2004). Periaqueductal Gray. Rat Nerv. Syst. 243–257 doi:10.1016/B978-012547638-6/50011-0.

67. Bandler, R. & Shipley, M. T. (1994). Columnar organization in the midbrain periaqueductal gray: modules for emotional expression? Trends in Neurosciences vol. 17 at 10.1016/0166-2236(94)90047-7.

68. Vaughn, E., Eichhorn, S., Jung, W., Zhuang, X. & Dulac, C. (2022). Three-dimensional Interrogation of Cell Types and Instinctive Behavior in the Periaqueductal Gray. bioRxiv.

69. Tovote, P., Esposito, M.S., Botta, P., Chaudun, F., Fadok, J.P., Markovic, M., Wolff, S.B., Ramakrishnan, C., Fenno, L., Deisseroth, K. et al. (2016). Midbrain circuits for defensive behaviour. Nature 534.

70. Souza, R. R. & Carobrez, A. P. (2016). Acquisition and expression of fear memories are distinctly modulated along the dorsolateral periaqueductal gray axis of rats exposed to predator odor. Behav. Brain Res. 315.

71. Mahn, M., Gibor, L., Patil, P., Cohen-Kashi Malina, K., Oring, S., Printz, Y., Levy, R., Lampl, I. Yizhar, O. (2018). High-efficiency optogenetic silencing with soma-targeted anion-conducting channelrhodopsins. Nat. Commun. 9.

72. Chen, T.W., Wardill, T.J., Sun, Y., Pulver, S.R., Renninger, S.L., Baohan, A., Schreiter, E.R., Kerr, R.A., Orger, M.B., Jayaraman, V. (2013). Ultrasensitive fluorescent proteins for imaging neuronal activity. Nature 499, 295–300.

73. Kabra, M., Robie, A. A., Rivera-Alba, M., Branson, S. & Branson, K. (2013). JAABA: Interactive machine learning for automatic annotation of animal behavior. Nat. Methods 10.

74. Berg, S., Kutra, D., Kroeger, T., Straehle, C.N., Kausler, B.X., Haubold, C., Schiegg, M., Ales, J., Beier, T., Rudy, M. et al. (2019). ilastik: interactive machine learning for (bio)image analysis. Nat. Methods 16.

75. Song, J.H., Choi, W., Song, Y.H., Kim, J.H., Jeong, D., Lee, S.H. and Paik, S.B. (2020). Precise Mapping of Single Neurons by Calibrated 3D Reconstruction of Brain Slices Reveals Topographic Projection in Mouse Visual Cortex. Cell Rep. 31.

76. Shang, C., Liu, Z., Chen, Z., Shi, Y., Wang, Q., Liu, S Cao, P. (2015). A parvalbumin-positive excitatory visual pathway to trigger fear responses in mice. Science (80-.). 348, 1472–1477.

77. Ahmadlou, M. & Heimel, J. A. (2015). Preference for concentric orientations in the mouse superior colliculus. Nat. Commun. 6.

78. Sommeijer, J. P., Ahmadlou, M., Saiepour, M. H., Seignette, K., Min, R., Heimel, J. A., & Levelt, C. N. (2017). Thalamic inhibition regulates critical-period plasticity in visual cortex and thalamus. Nat. Neurosci. 20, 1716–1721.

79. Ahmadlou, M., Zweifel, L. S. & Heimel, J. A. (2018). Functional modulation of primary visual cortex by the superior colliculus in the mouse. Nat. Commun. 9.

80. Montijn, J.S., Meijer, G. T., & Heimel, J. A. (2023). A new family of statistical tests for neural spiking and autocorrelated timeseries data. bioRxiv 1–38.

