## Supplemental Figures for "Cell Type-specific Hypothalamic Pathways to Brainstem Drive Context-dependent Strategies in Response to Stressors"

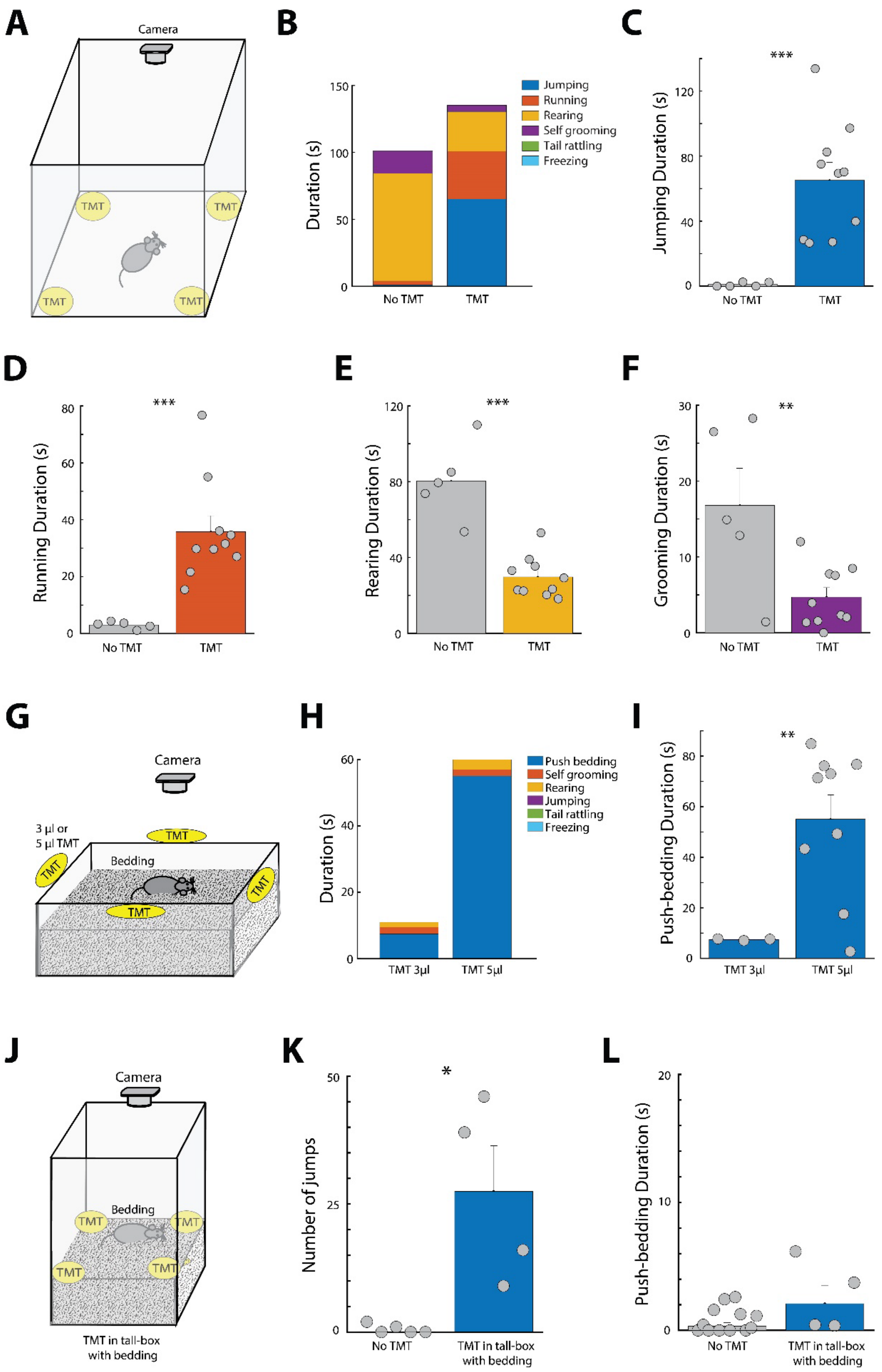

A
Camera
TMT
B
Duration (s)
Jumping
Running
Rearing
Self grooming
Tail rattling
Freezing
No TMT
TMT
C
Jumping Duration (s)
***
D
Running Duration (s)
E
Rearing Duration (s)
F
Grooming Duration (s)
**
G
3 μl or
5 μl TMT
Bedding
H
Push bedding
Self grooming
Rearing
Jumping
Tail rattling
Freezing
TMT 3μl
TMT 5μl
I
Push-bedding Duration (s)
J
TMT in tall-box
with bedding
K
Number of jumps
*
L
Push-bedding Duration (s)

**Figure S1. Behavior of mice in the inside- and outside-stressor tests. Related to Figure 1.**

(**A**) Schematics of an inside-stressor test with TMT cottons in all corners. (**B**) Stacked bar graph shows duration for each action (running, jumping, rearing, self-grooming, rearing, tail rattling and freezing) in the inside-stressor test taken by C57BL/6 mice in Figure 1B. Bar graphs show duration of jumping (**C**), running (**D**), rearing (**E**), and self-grooming (**F**) in the inside-stressor test. (**G**) Schematics of an outside-stressor test with 3 µl or 5 µl TMT cottons sticking at all edges outside the home-like cage with a thick layer of beddings. (**H**) Stacked bar graph shows duration for each action (push-bedding, self-grooming, rearing, jumping, tail rattling and freezing) in the outside-stressor test (with 3 µl or 5 µl TMT cottons) taken by C57BL/6 mice. (**I**) The bar graph shows duration of push-bedding in the outside-stressor test with 3 µl TMT cottons (N = 3 mice) and with 5 µl TMT cottons (N = 9 mice) taken by the mice in (H). (**J**) Schematics of an inside-stressor test in a tall box with beddings and TMT cottons located at all corners. (**K**) Bar graph shows the number of jumps taken by C57BL/6 mice. (**L**) Bar graph shows duration of push-bedding. *: $p < 0.05$, **: $p < 0.01$, ***: $p < 0.001$.

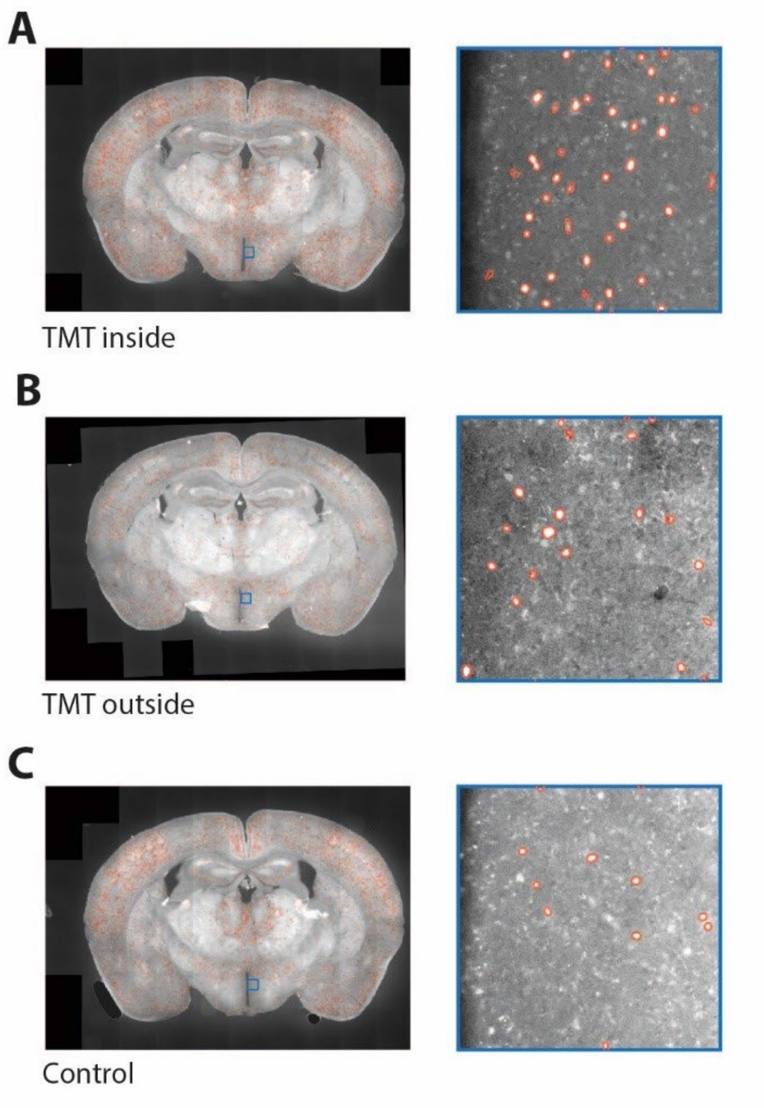


**Figure S2. DMH neurons are activated in the inside- and outside-stressor tests. Related to Figure 2.**

(**A**) The left shows an example coronal slice of mouse that underwent the TMT inside paradigm, stained for c-Fos. In red the cells detected by Ilastik are shown. The right shows an enlargement of the part of the DMH shown indicated by the blue square. **(B)** Same as A for an animal that was underwent the TMT inside paradigm. **(C)** Same as A for an animal that was placed as animals in the TMT inside paradigm but that was not exposed to TMT.

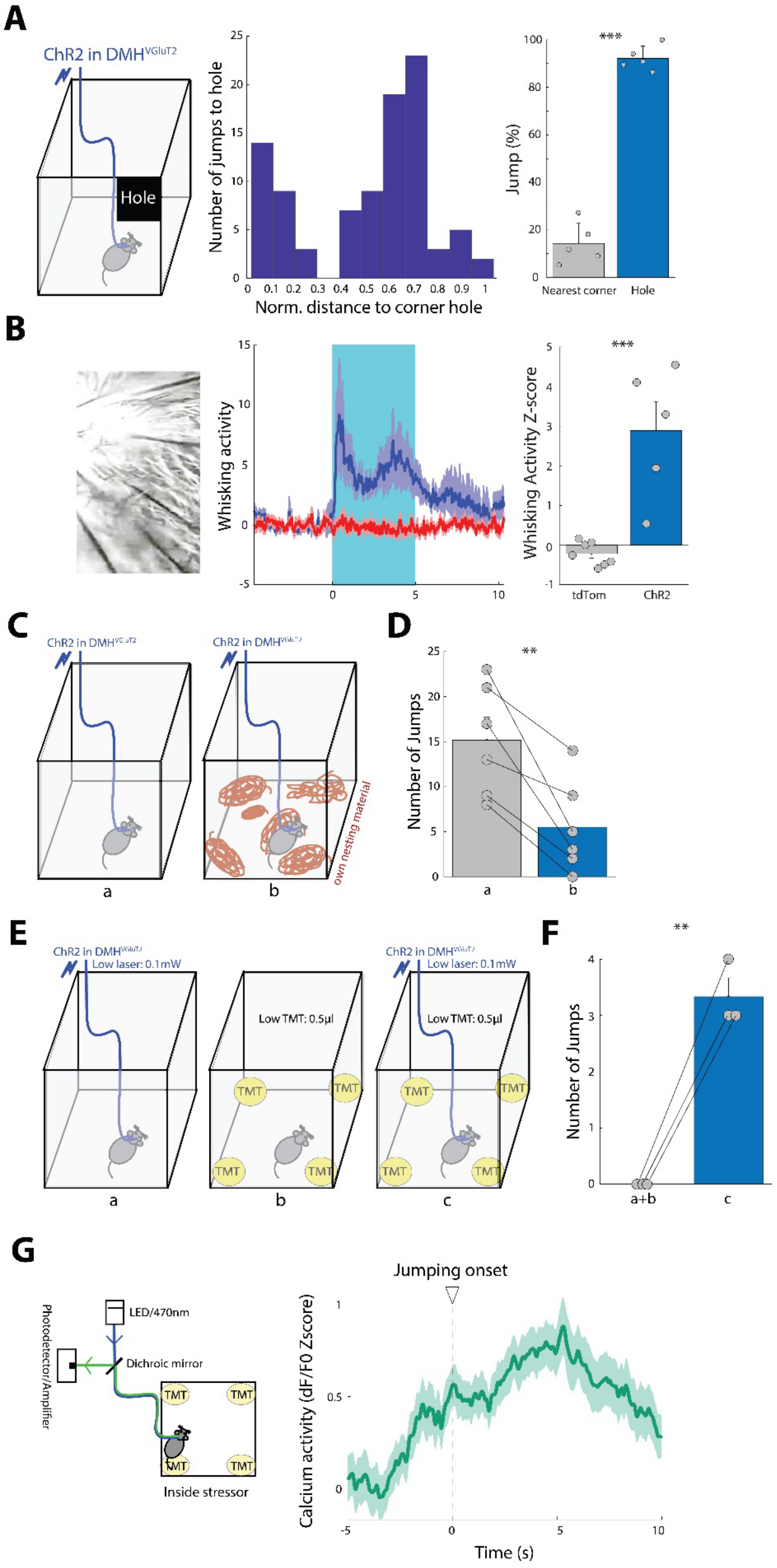
A
ChR2 in DMH$^{VGluT2}$
Hole
Number of jumps to hole
Norm. distance to corner hole
Jump (%)
***
Nearest corner
Hole
B
Whisking activity
Whisking Activity Z-score
***
tdTom
ChR2
C
ChR2 in DMH$^{VGluT2}$
own nesting material
a
b
D
Number of Jumps
**
E
ChR2 in DMH$^{VGluT2}$
Low laser: 0.1mW
Low TMT: 0.5µl
TMT
a
b
c
F
Number of Jumps
**
a+b
c
G
Photodetector/Amplifier
LED/470nm
Dichroic mirror
Inside stressor
Jumping onset
Calcium activity (dF/F0 Zscore)
Time (s)

**Figure S3. DMH$^{VGluT2}$ neurons modulate arousal level and goal-directed jumping. Related to Figure 2.**

(**A**) Schematics of photo stimulation of DMH$^{VGluT2}$ neurons in the box (without TMT) with a hole in one top corner (corner hole). Histogram of number of jumps to the corner hole as a function of normalized distance (at the start of the photo stimulation) to the corner hole (N = 5 mice). The bar graph shows the number of jumps to the nearest top corner and the corner hole after each photo stimulation normalized to the number of photo stimulation trials (in percentage). (**B**) Z-score (middle) and mean Z-score (right) of whisker activity of tdTom (red; N = 7 mice) and ChR2 (blue; N = 5 mice) mice with photo stimulation from 0 to 5 s. (**C**) Schematics of photo stimulation (laser level: 3 mW) of DMH$^{VGluT2}$ neurons in the box with two stress levels: a: without and b: with their own nesting material. (**D**) The bar graph shows the number of jumps in the ChR2 injected mice in both conditions in (A) (N = 6 mice). (**E**) Schematics of photo stimulation of DMH$^{VGluT2}$ neurons in the box with low laser level (0.1 mW) without low TMT (0.5 µl) (a), with low TMT and without photo stimulation (b) and with both low laser level and low TMT (c). (**F**) The bar graph shows the number of jumps in the ChR2 injected mice in 'a' together with 'b' conditions and in 'c' condition in (C) (N = 3 mice). (**G**) Left: schematic of photometry recording from DMH$^{VGluT2}$ neurons during inside-stressor situation. Right: calcium activity of DMH$^{VGluT2}$ neurons (dF/F0 Z-score) from the mice shown in Figure 2P in the inside-stressor test, averaged over trials with jumping responses (20 trials). The light color around the average indicates the standard error. Time 0 s is the onset of jumping. **: $p < 0.01$, ***: $p < 0.001$.

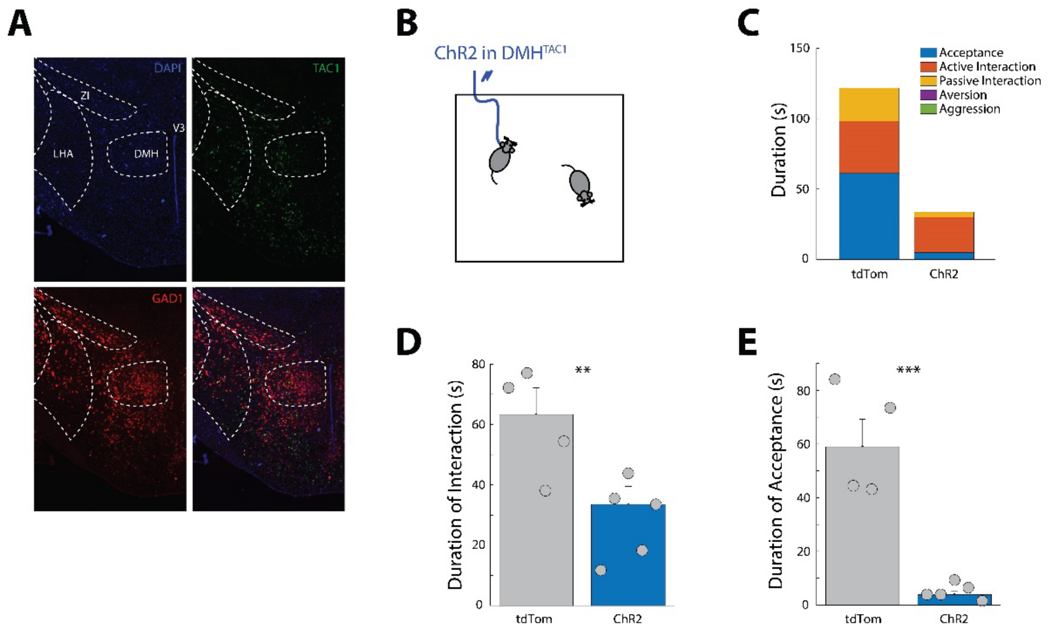


**Figure S4. DMH$^{TAC1}$ neurons are inhibitory and modulate social behaviors. Related to Figure 3.**

(**A**) Experiment #180304064 of Allen Mouse Brain Connectivity Atlas (https://connectivity.brain-map.org/transgenic/experiment/180304064): DAPI staining (blue), expression of tdTomato in DMH neurons in a TAC1-Cre crossed with tdTomato reporter mouse line (Ai14) (green) and mRNA in-situ hybridization of GAD1 expressing neurons in DMH. ZI: Zona Incerta, LHA: Lateral Hypothalamic Area, DMH: Dorsomedial Hypothalamus, V3: Third Ventricle. (**B**) Schematics of photo stimulation of DMH$^{TAC1}$ neurons during social interaction with a novel conspecific. (**C**) Stacked bar graph shows duration for each action in the social interaction test in tdTomato and ChR2 expressing groups (N = 4 and 5 mice, respectively). Acceptance: when the resident mouse accepts to stay (without taking any investigatory interaction), while it is investigated by the intruder mouse. Active interaction: when the resident mouse actively approaches the intruder mouse initiates the interaction and investigates the intruder mouse. Passive interaction: when the interaction of the resident mouse with the intruder mouse starts with approach and initiation of the intruder mouse. Aversion: when the resident mouse shows aversive behavior in response to the intruder mouse, such as running away and tail rattling. Aggression: when the resident mouse shows aggressive behaviors such as fighting, boxing, wrestling and biting. (**D**) The bar graph shows duration of interaction (passive + active) in the social interaction test taken by the mice in (C). (**E**) The bar graph shows duration of acceptance in the social interaction test taken by the mice in (C). **: $p < 0.01$, ***: $p < 0.001$.
